# PAIRWISE: Deep Learning-based Prediction of Effective Personalized Drug Combinations in Cancer

**DOI:** 10.1101/2024.11.04.621884

**Authors:** Chengqi Xu, Ilkay Us, Jake Cohen-Setton, Marta Milo, Ben Sidders, Jude Fitzgibbon, Ari M. Melnick, Heng Pan, Olivier Elemento, Krishna C. Bulusu

## Abstract

Combination therapies offer promise for improving cancer treatment efficacy and preventing recurrence. However, identifying optimal drug combinations tailored to specific cancer subtypes and individual patients is extremely challenging due to the vast number of possible combinations and tumor heterogeneity. To address this gap, we take a machine learning approach combining deep learning with transfer learning to incorporate prior scientific knowledge and predict drug synergy based on tumor-specific transcriptome profiles. This approach, called PAIRWISE, explicitly modeled synergistic effects of drug combinations in cancer cell lines or individual tumor samples based on drug chemical structures, drug targets, and transcriptomes of inferred samples. PAIRWISE outperformed competing models with an area under the receiver operating characteristic curve (AUROC) of 0.85 on held-out cancer cell lines. When applied to an independent dataset of combinations with Bruton Tyrosine Kinase inhibitors (BTKi) in Diffuse Large B Cell Lymphoma (DLBCL) cell lines, PAIRWISE accurately predicted synergistic drug combinations with an AUROC of 0.72. To further confirm the robustness of PAIRWISE predictions, we performed an in silico, patient profile-directed screen for other compounds that would synergize with BTKi in DLBCL patients, and confirmed the synergy of the predictions using a panel of eight non-Hodgkin lymphoma cell lines. These findings demonstrate the ability of PAIRWISE to nominate effective personalized drug combinations, accelerating the development of precision oncology.

## Introduction

Combination therapies using multiple anti-cancer drugs together have become a cornerstone of cancer treatment^1^. Regimens combining chemotherapy, targeted agents, and immunotherapies have improved cancer outcomes in the last decades^2–5^. However, identifying optimal drug combinations for each cancer type and individual patient remains challenging. The search space of possible combinations is vast, and responses can vary greatly due to tumor heterogeneity. High-throughput screening (HTS) provides an unbiased approach to testing large numbers of combinations, but usually lacks patient-derived cell models and cannot identify patient-specific combinations^2,6–9^. Current empirical methods for designing combinations also have limitations in predicting patient-specific synergistic regimens. There is thus an unmet need for accurately predicting personalized synergistic drug combinations.

Advanced computational methods could greatly reduce the search space and minimize the experimental efforts required to determine effective drug combinations. ^10^Traditional machine learning (ML) models could learn functional mappings between high-dimensional input data and drug synergy measured by the Loewe or Bliss additivity model^10^, requiring extensive feature selection before training^11–14^. Recent deep learning (DL) models adopt a multimodal architecture^15–30^, bypassing the reliance on singular data modalities. However, extant models rarely use pre-trained techniques to represent inherent features of both chemical and sample-specific transcriptional modalities. It is challenging for them to accurately predict drug synergy in cancers with limited cases in the training set.

To address these gaps, we developed a DL model PAIRWISE, which employed a multimodal architecture to integrate different modalities influencing drug responses, including chemical structures, drug targets, and transcriptomes of individual tumor samples. Modalities were encoded with specialized neural networks and integrated using attention mechanisms that focus learning on relevant features. PAIRWISE also utilized transfer learning, wherein the model is pre-trained on large drug and transcriptome datasets, thus learning generalized representations of drug chemical structures and sample-specific transcriptomes.

We first applied PAIRWISE and benchmarked methods to a curated drug combination screening dataset collected from 13 public sources across hematological malignancies and solid tumors^31^. The curated dataset was named the p13 dataset and has ∼200,000 screened drug combinations across 167 cell lines. Compared with benchmarked methods, PAIRWISE achieved better performance across several metrics including AUROC (the area under the receiver operating characteristic curve) and AUPRC (the area under the precision-recall curve) when drugs, drug combinations, or cancer cell lines were held out. Moreover, PAIRWISE showed high sensitivity and specificity for predicting synergistic drug combinations containing Bruton Tyrosine Kinase inhibitors (BTKi) in an independent dataset^32^. Here, we experimentally demonstrated that several of the novel synergistic BTKi combinations are indeed synergistic in Diffuse Large B Cell Lymphoma (DLBCL) cell lines. Combinations of BTKi with agents targeting DNA Damage Response (DDR) pathways demonstrated marked synergy in aggressive subtypes of DLBCL. Finally, PAIRWISE was applied to transcriptome profiles of primary patient DLBCL tumors (LS cohort) and predicted candidate drugs with synergy with BTKi, such as PARP and DNA-PK inhibitors. These predictions suggest potential therapeutic strategies targeting DNA damage response (DDR) pathways in aggressive DLBCL subtypes, providing valuable insights for personalized treatment options^33^ PAIRWISE represents a novel DL framework for identifying optimal drug combinations tailored to specific cancer subtypes and individual patients, paving the way toward an enhanced understanding of the role of personalized drug combinations in cancer.

## Results

### PAIRWISE is a novel DL model for personalized drug combination prediction

To predict patient-specific effective drug combinations, we developed PAIRWISE - an integrative DL approach that considers both the mechanism of drug combination and transcriptome of individual cancer samples. PAIRWISE can predict whether two compounds or drugs are likely to achieve synergistic cell killing in individual cancer samples, even for drugs that are not present in the combination screening data. To fulfill this objective, PAIRWISE uses transfer learning to first ground the model in general domain knowledge before fine-tuning it on our curated large-scale combination screening data. Also, a multimodal architecture allows PAIRWISE to learn higher-order relationships among drug chemical structures, drug target interactions (DTIs), and sample-specific transcriptome.

To be specific, PAIRWISE takes three sets of features as input: a combination of two drug chemical structures, a combination of two DTIs, and the transcriptome profile of an individual sample (**Fig. 1a**). PAIRWISE outputs the binary prediction of the combination effect (synergistic or not) of two drugs in the inferred sample. To create vectorized input features, we represented drug chemical structures using the Simplified Molecular Input Line Entry System (SMILES)^34,35^. SMILES were then converted into molecular graphs encoded as adjacency matrices, in which each node represents an atom and each edge represents a chemical bond between atoms. DTIs were converted into binary vectors recording whether a single gene is the drug target (1 indicates yes while 0 indicates no) across all known targets of at least one drug (*n* = 4,098) in Drug Target Commons (DTCs)^36^. Besides primary and secondary targets, DTCs also contains indirect targets related to disease or drug response, hence could capture the whole spectrum of potential targets. Transcriptome profiles of individual samples were represented as an expression matrix of all genes. Vectorized features were then fed into embedding layers of PAIRWISE. Embedding layers serve as feature extractors and consist of three branches: (1) the chemical fingerprint branch, (2) the DTI branch, and (3) the cancer sample branch (**Fig. 1b**). The chemical fingerprint branch leverages transfer learning techniques built upon a chemical pre-trained foundation model (PFM)^37^. This general model was pre-trained on 11 million chemical compounds, which can enable reasonable parameter initiation and easy fine-tuning. Compared with simple chemical descriptors such as SMILES or Morgan fingerprints^38^, chemical PFMs provide more sophisticated representations of atoms and chemical bonds. Parameters of the chemical PFM were updated and used to guide PAIRWISE after fine-tuning with the p13 dataset. The parameters update was frozen in the test dataset. The DTI branch was implemented with a convolutional neural network (CNN), which condensed 4,098 drug targets into 500 features that contain critical information about DTI. In the cancer sample branch, PAIRWISE represented transcriptomes of individual samples using an auto-encoder pre-trained on *in vitro* Cancer Cell Line Encyclopedia (CCLE) cell-line samples^39^ and clinical The Cancer Genome Atlas (TCGA) patient samples^40^ with corresponding transcriptomes profiles. Instead of the raw expression matrix, the pre-trained autoencoder learns shared biological signals between datasets common to cell-line and patient data. In this way, the knowledge learned from the training samples can be transferred to a new sample. Next, an attention-based feature fusion module was used to combine features in each branch (**Fig. 1c**). In the attention layer, each feature embedding was applied with a scaled dot product-based attention with the embedding of other modalities to produce an update feature embedding. The five resulting embeddings were combined across all three branches in a single-layer neuron to produce joint features, which were then fed into an output layer to generate the binary prediction of synergy for a particular drug combination in a given sample (drug combinations with a probability > 0.5 were classified as synergistic) (**Fig. 1d**).

**Fig. 1:**
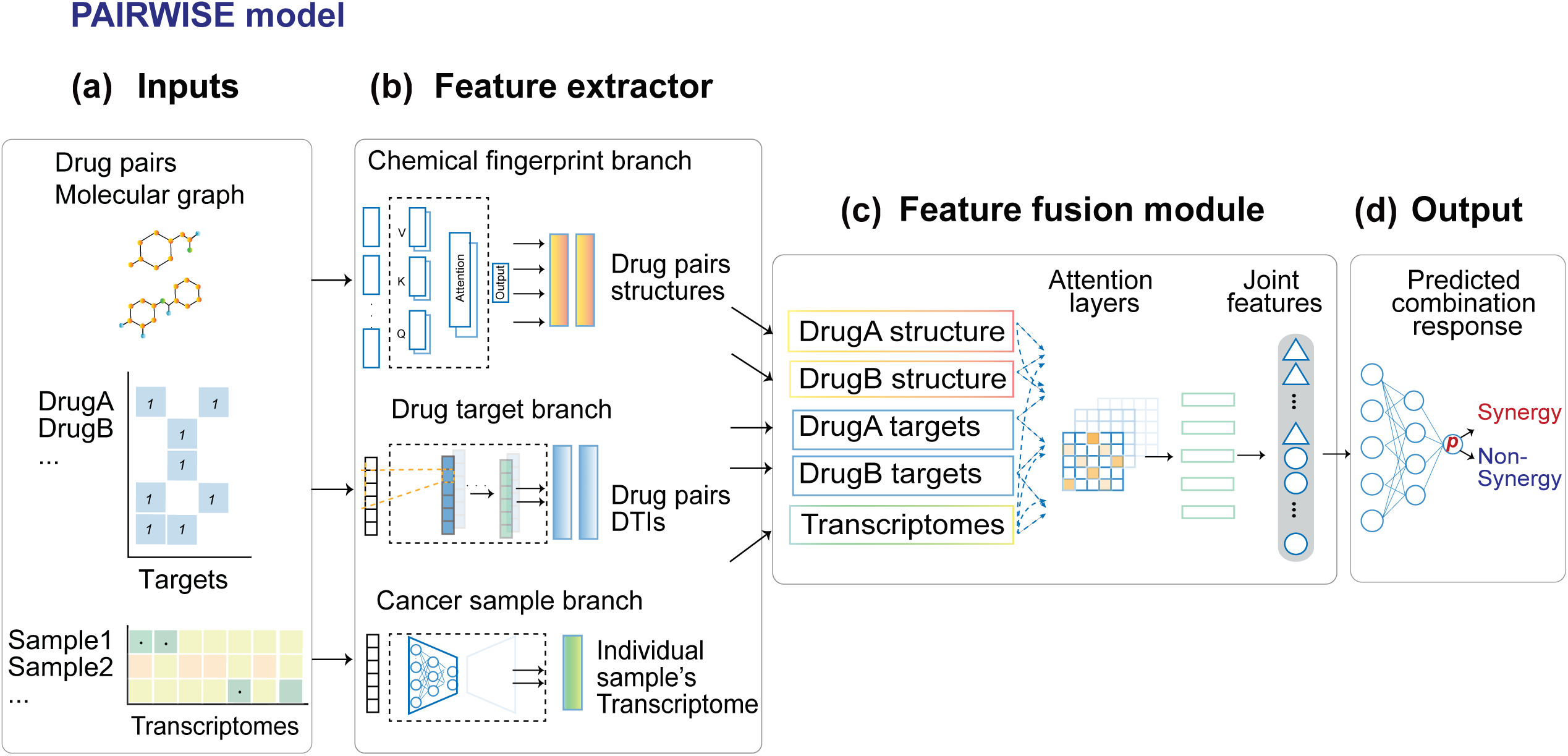
The PAIRWISE model for predicting drug synergy. **(a)** PAIRWISE takes three inputs: the drug combination molecular graphs representing chemical information as adjacency matrices, which capture the atoms and chemical bonds of the input compounds structures. The drug target is generated as an equal-length vector recording the binary status whether that input drug has a specific drug target. The cancer sample used transcriptome values as input, which denote the individualized transcriptome profile of that sample. **(b)** In feature extractor module, for the chemical fingerprint branch, a chemical foundation model is trained on 10 millions of molecule structures, and it is fine-tuned to guide the drug pairs structures optimization; for the drug target branch **(c)** In the feature fusion module, two vectors representing drug structures embeddings, two vectors representing drug targets embedding, and a single vector representing cancer sample with the same length are aligned and updated in the attention layer. The updated embeddings from an attention mechanism are linearized into a joint feature embedding vector. **(d)** In the output layer, the join feature embedding is passed into a multi-layer perceptron to produce the predicted synergy probabilities, and consequently measure a given drug combination effect between synergy and non-synergy status of an individualized cancer sample.

### PAIRWISE outperforms state-of-the-art models across cancer cell lines from different tissues

To optimize the parameters of PAIRWISE, we first aggregated 13 public HTS assays of drug combinations using unique criteria (**Methods**; **Supplementary Table. 1**). Synonyms of each compound were determined by the Tanimoto similarity of Morgan fingerprints and were assigned with unique DrugBank IDs (**Supplementary Fig. 1**). We determined using Morgan fingerprints since target similarities are quite low (**Supplementary Fig. 2)**. Cell lines and synonyms were assigned unique DepMap IDs using a name-mapping table derived from the DepMap portal^39^ (**Supplementary Fig. 1**). Synergy scores of drug combinations were calculated based on dose-response metrics, including Bliss, HSA, Loewe, and ZIP. This curated dataset (p13) contained 279,452 unique drug combinations, 2,614 drugs, and 167 cancer cell lines. We also implemented an end-to-end pipeline to assist in the development and evaluation of drug combination prediction models (Supplementary Fig. 2).

To evaluate the performance of PAIRWISE, we tested our model through 3 distinct settings and benchmarked five published DL models (**Supplementary Table. 2**). In each setting, the p13 dataset was equally split into 5 folds, which do not share common drug combinations (leave-combo-out), drugs (leave-drug-out), or cancer samples (leave-sample-out) (**Table 1**). Specifically, we employ a 5-fold cross-validation within each setting, where the training set is further divided into training and validation subsets using a 90/10 ratio. These settings were used to evaluate the ability of models to generalize from the training data to unseen drug combinations, individual drugs, or cancer samples. In the leave-sample-out setting, 167 cell lines were divided into 4 clusters based on the expression of the top 1,000 most variable genes used an affinity propagation method. The biggest cluster with 93 cell lines was used as the training set, and the remaining cell lines were used as the test set. PAIRWISE showed superior performance compared to other models when assessed using AUROC and AUPRC (the area under the precision-recall curve) in all three settings (**Fig. 2a-b**). Before being converted to binary classifications, PAIRWISE and benchmarked methods outputted continuous probabilities of drug synergy. PAIRWISE also achieved the highest Spearman correlation between predicted synergy probabilities and experimental synergy scores in all settings (**Fig. 2c**; **Table 2**).

**Fig. 2:**
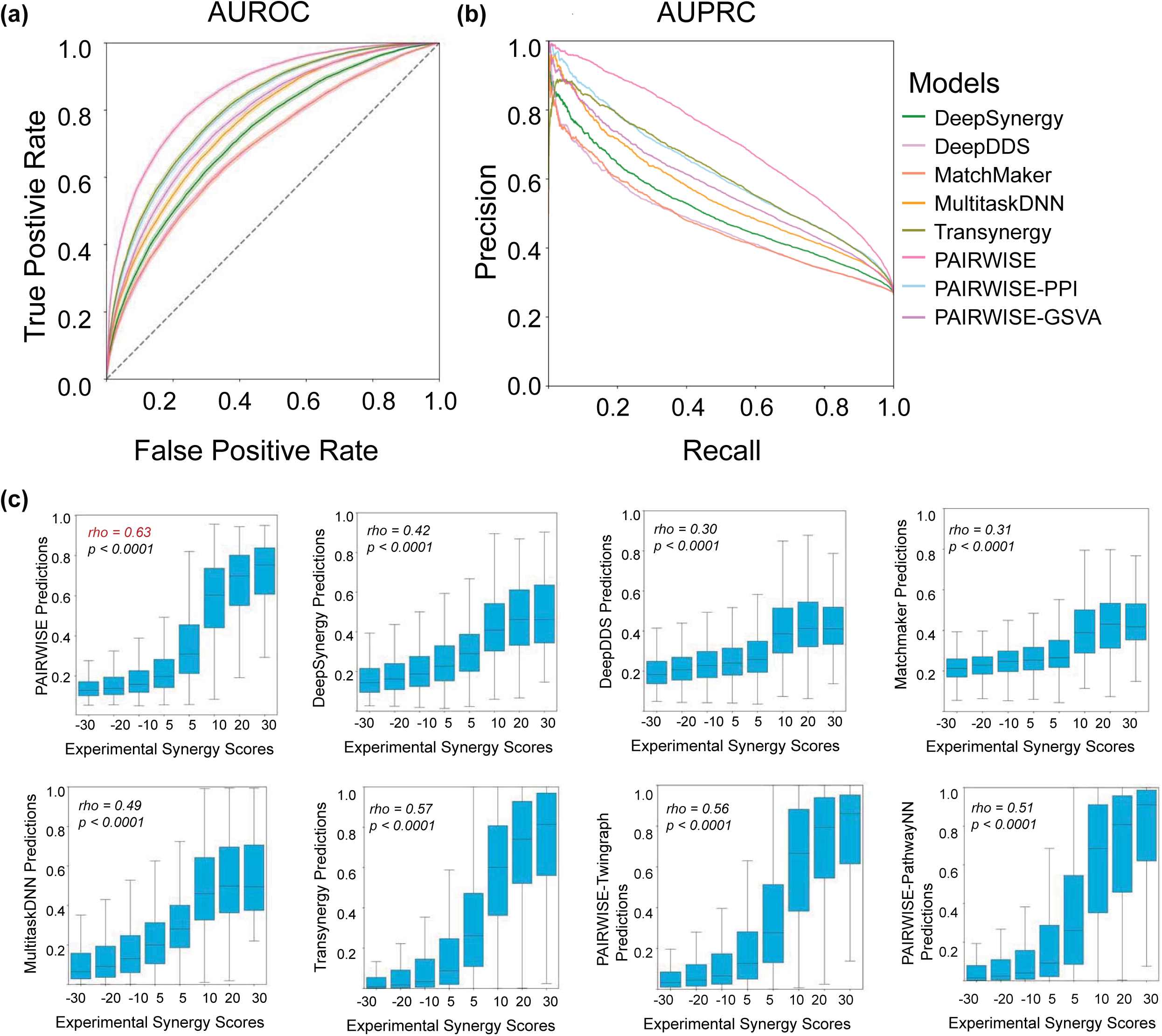
Predictive performance. (a-b) The performance of the suite of predictive models for drug synergy across metrics. For each model, which predicts drug synergy as defined by each respective model, the performance was measured by AUROC and AUPRC. **(c)** Predicted versus actual drug combination synergy scores across all (cell line, drug combinations) tuples studied. Box plots show each bin’s 25th, 50th, and 75th percentiles; whiskers show maximum and minimum values.

To assess PAIRWISE performance in diverse tissues, we explored the performance of the model in 10 different cancer types of the p13 test dataset. The AUROC showed consistent robust performance (0.858±0.003), including cancers in colon, breast, haematopoietic and lymphoid, etc. (**Supplementary Fig. 3)**. By contrast, we found that the published model (Multitask-DNN), which specifically designed for transfer learning in tissues with limited training data such as bone and prostate cell lines, has limited performance (**Supplementary Table. 3**). PAIRWISE predictions exhibited strong correlations (Spearman’s Coefficient, 0.508-0.606) with experimental synergy scores across different tissues (**Supplementary Fig. 3**). Compared with other published models, PAIRWISE exhibited consistent AUROC performance across tissues, including cancers with limited training data (**Supplementary Fig. 4**). We conclude that PAIRWISE learns cross-tissue generalizable patterns of drug combinations, and provides accurate prediction of combination treatments.

### PAIRWISE benefits from the branch representation methods in drug synergy prediction

To systematically evaluate the importance of each of the three input and representation branches in PAIRWISE, we replaced individual representation methods with alternatives. Six PAIRWISE models were created, and their predictive performance was compared to PAIRWISE on the p13 dataset. For the chemical fingerprint branch, we replace the Chemical PFM with SMILES (PAIRWISE-SMILES) or Morgan (PAIRWISE-Morgan) Fingerprints to represent a drug compound chemical structure. For the drug target branch, we replaced DTCs with DrugBank^41^ (PAIRWISE-DrugBank), a widely used database, as well as an alternative approach to propagate DrugBank with PPI (PAIRWISE-PropagatedDrugbank). For the cancer sample branch, we introduced two different cancer sample representations based on PPI and Gene set variation analysis (GSVA) scores to surrogate pre-training techniques (**Methods; Supplementary Fig. 5**). Compared with all the variant models, PAIRWISE had the best synergy prediction accuracy on the p13 dataset (**Table 3**). PAIRWISE increased performance by correctly classifying more synergy cases when compared to PAIRWISE-DrugBank and PAIRWISE-PropagatedDrugBank (**Supplementary Fig. 6**). Replacing pre-trained TCGA encoders with PPI and GSVA decreases the performance of synergy prediction. Of note, the AUPRC of PAIRWISE-SMILES and PAIRWISE-Morgan were the lowest among all the models, suggesting the chemical finger branch featured with chemical PFM may have the most significant contribution to the prediction accuracy in PAIRWISE.

### PAIRWISE accurately predicts drug synergy in an independent DLBCL HTS dataset

Next, we sought to assess the performance of PAIRWISE in an independent drug combination dataset. We used an HTS dataset testing the efficacy and synergy of a BTK inhibitor in DLBCL^32^ (**Fig. 3a**). The HTS study analyzed 466 combinations of ibrutinib with approved or investigational drugs in the TMD8 DLBCL cell line. Drugs without known chemical structures and gene targets were filtered out, and 214 drugs were left for validation. On this dataset PAIRWISE achieved a robust prediction accuracy of synergistic pairs (AUROC = 0.720±0.003, **Fig. 3b-c**). Of note, this task is very challenging since there were only a small number of records from 3 hematopoietic cell lines in the training cohort (*n* = 2,445 out of 279,452; 1% of the training cohort), and no records were from TMD8. Combinations that were predicted to have synergy showed significantly higher synergy in the experimental combinatorial viability assay, compared with those that were predicted to be only non-synergistic (*P* = 0.0014, Wilcoxon rank sum test). These results suggest that the model is still effective with cell lines or drug compounds that were never seen before by the model.

**Fig. 3:**
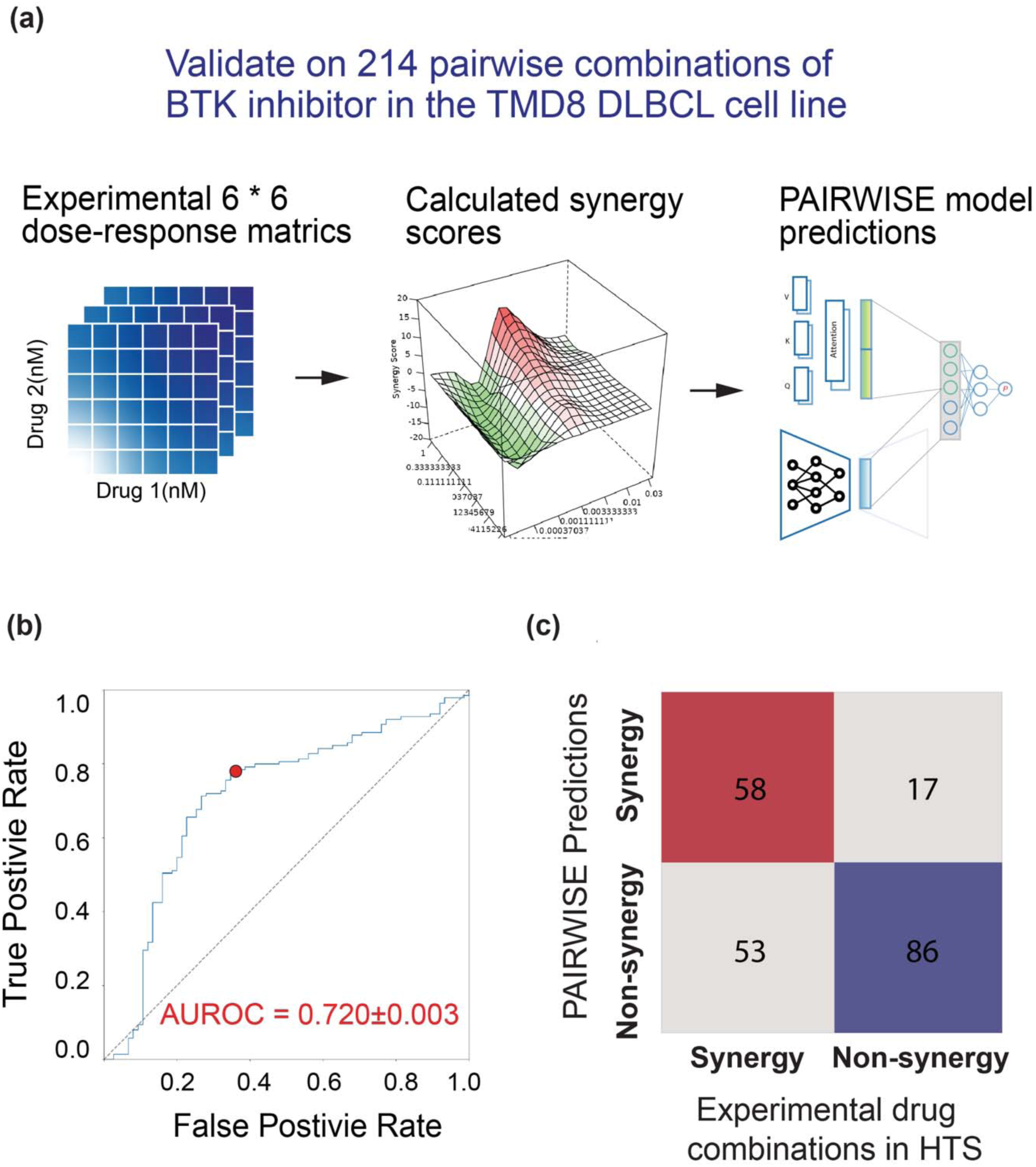
PAIRWISE predicts accurately combinatorial responses with the external experimental DLBCL HTS design. **(a)** Flowchart of the analysis procedure. **(b)** AUROC of PAIRWISE performance in distinguishing effective from ineffective drug combinations. (**c)** Confusion matrix for point indicated in (b) demonstrating the best performance of PAIRWISE against the DLBCL HTS dataset.

### High-throughput screening validates synergistic BTKi combinations recommended by PAIRWISE in DLBCL

One of the strengths of PAIRWISE is that despite being trained on cell lines, it can in theory be applied to primary patient samples. Encouraged by our results in DLBCL, we thus sought to predict drug combination synergy in 562 primary DLBCL patient samples (LS cohort) with a median age of 64 years old (range, 25–74) where transcriptome profiles were available^33^. Most patients presented relatively high-risk disease: 45% with Ann Arbor stage III or IV, and 70% with IPI group intermediate or high. We continued to focus on combinations involving BTKi since there is continued interest in using them in DLBCL. PAIRWISE was applied to predict combination synergy of BTKi in conjunction with 211 drugs from NCATS MIPE library (a resource containing oncology-focused and clinic-prioritized targeted therapies) and 57 additional investigational drugs (**Fig. 4a**). In total, 30 out of 268 combinations were identified as likely synergistic based on the rank of average predicted synergy scores across all the samples (**Fig. 4b**). Predicted synergistic combinations included drugs that play critical roles in DNA repair, epigenetic modification, JAK/STAT signaling, and PI3K/AKT signaling (**Supplementary Table. 4**).

**Fig. 4:**
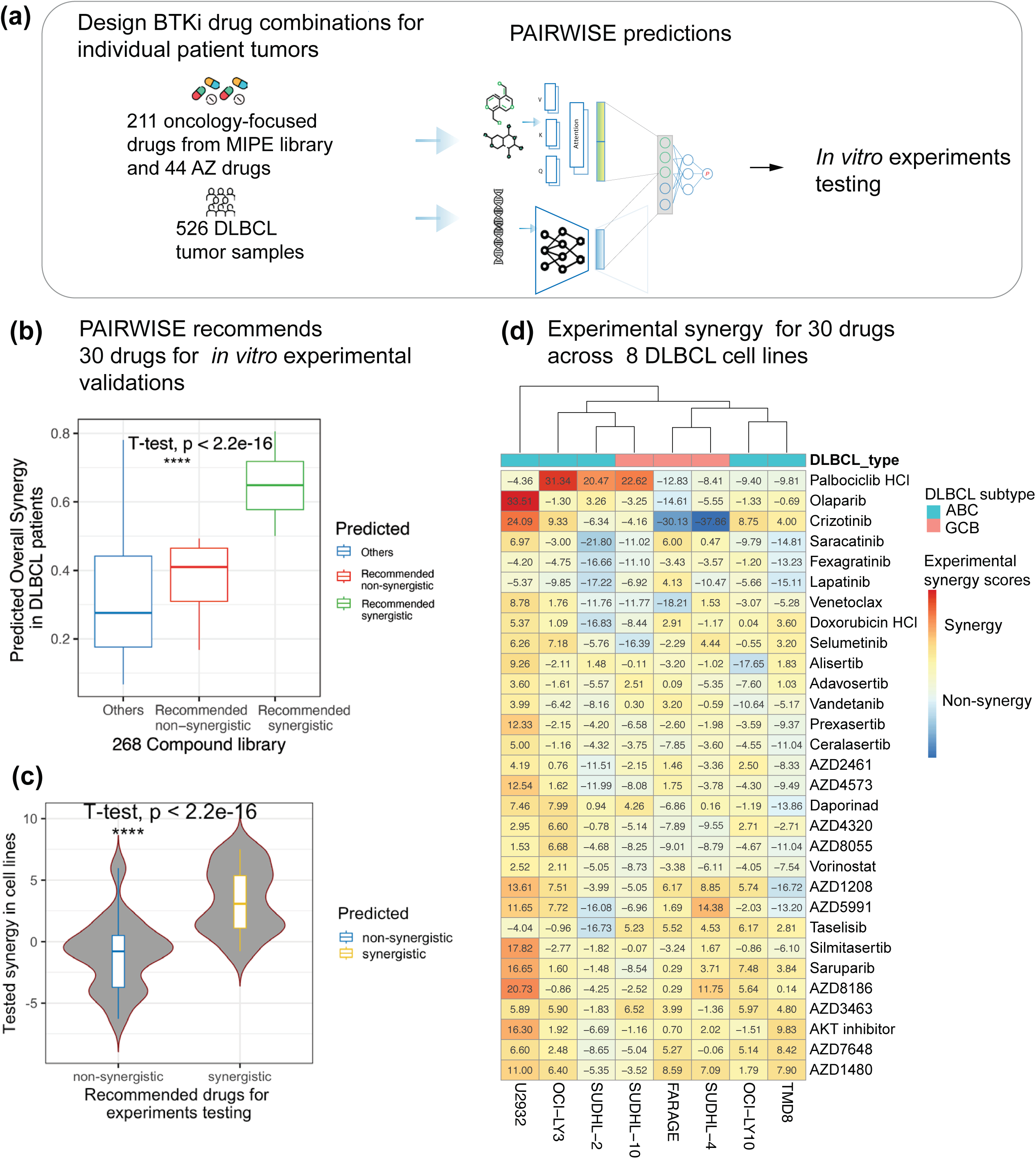
Experimental validation of top predicted combinations. **(a)** Flowchart of the analysis procedure. **(b)** PAIRWISE model recommends 30 drugs with predicted synergy from 211 pairwise combinations with secondary drugs from diverse target classes from MIPE library and 44 AZ drugs. **(c)** Experimental validation, quantified by Loewe synergy scores. **(d)**. Distribution of combination scores of each tested drug combination.

We sought to confirm their synergy by testing 30 predicted drug combinations in 8 DLBCL cell lines in 6 * 6 dose-response combination metrics to (**Methods; Fig. 4d**). We found that synergistic drug combinations nominated by PAIRWISE were strongly and significantly enriched for synergistic cell killing outcomes, in contrast to combinations predicted to be non-synergistic combinations (**Fig. 4c**). As expected, the experimental synergies were heterogenous in different ABC and GCB DLBCL cell lines (**Fig. 5a-b**). Notably, we observed a high concordance between median predicted synergies in DLBCL patients and median experimental synergy scores in DLBCL cell lines across predicted drug combinations (Pearson’s Coefficient 0.55, *P <* 0.001, **Fig. 5c)**. This correlation was consistent in GCB and ABC DLBCLs (Pearson’s Coefficient 0.54, *P <* 0.001, **Fig. 5d**; Pearson’s Coefficient 0.62, *P <* 0.001, **Fig. 5e)**. These observations confirm the ability of PAIRWISE to robustly predict drug synergy in different DLBCL subtypes. PAIRWISE identifies the combination of olaparib and saruparib, PARP inhibitors, with acalabrutinib as having more synergy in more aggressive ABC DLBCL. The capacity of PARP inhibitors to synergize with acalabrutinib, specifically by targeting and overcoming single drug resistances has recently been demonstrated for mantel cell lymphoma^42^. Interestingly, PAIRWISE also identifies a combination with Acalabrutinib for which synergy is also associated with alterations in DDR pathway, such as the combination of DNA-PK inhibitor AZD7648. Recent studies established a connection between DDR activation and BCL-2 overexpression, serving as the biological underpinning for the unfavorable prognosis and chemoresistance evident in DLBCL subtypes^43^. PAIRWISE identifies a number of additional drugs that may be combined with Acalabrutinib to the same effect, including the AKT inhibitor capivasertib, the multitargeted tyrosine kinase inhibitor crizotinib and cyclin-dependent kinase 4/6 inhibitor palbociclib.

**Fig. 5:**
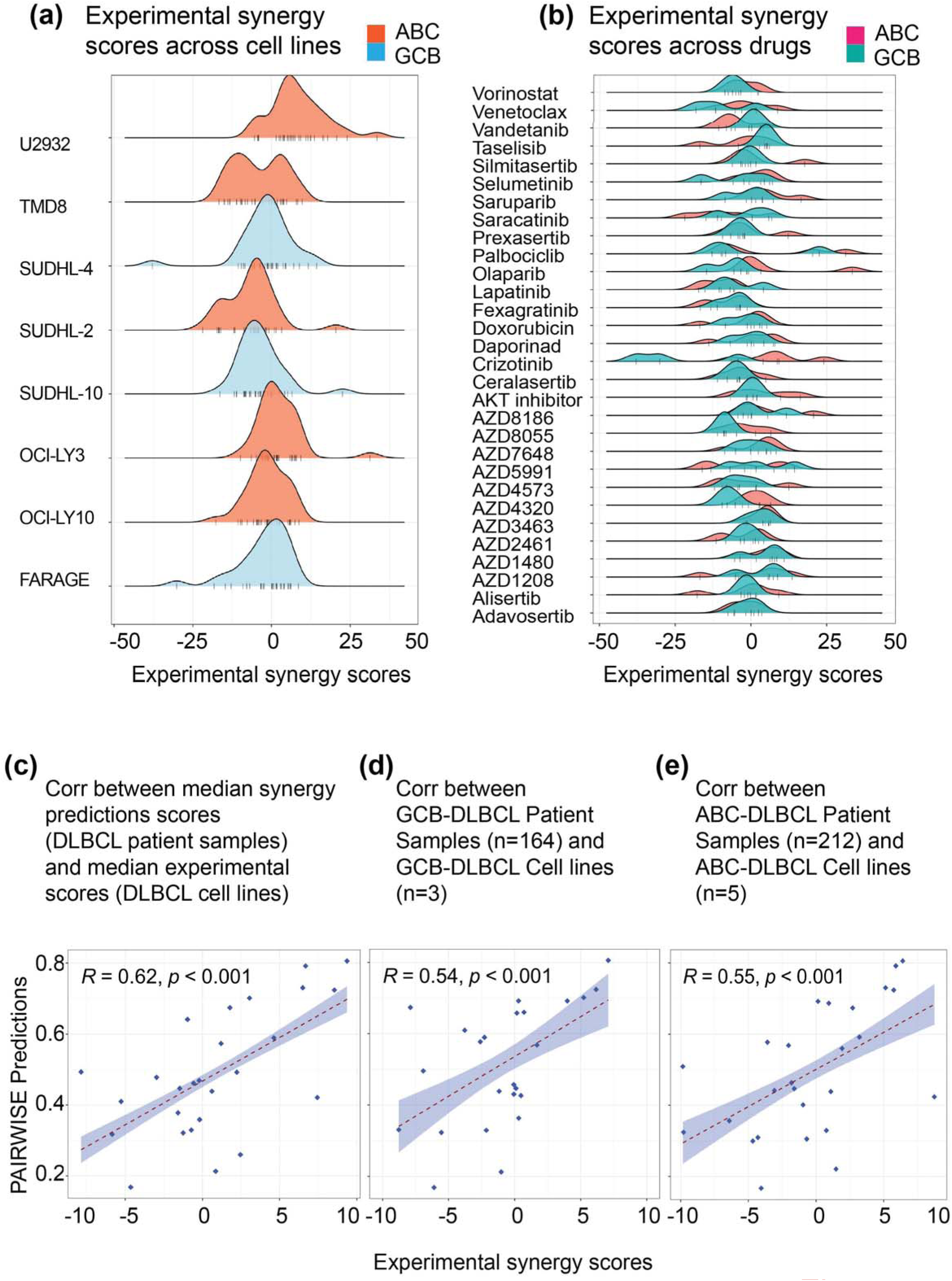
Experimental validation of top predicted combinations (a-b) Distribution of synergy scores across cell lines/ drugs. **(c-e)** Concordance between median synergy predictions in DLBCL patients and median measured synergy scores in DLBCL cell lines across recommended drugs.The p-value indicates significance by Pearson correlation.

### PAIRWISE-based combination stratification among patients with DLBCL

Finally, we sought to evaluate whether PAIRWISE could be used to stratify cancer patients into responsive and non-responsive populations to drug combinations. We analyzed the LS cohort in more detail after seeing the model had been validated. Out of 562 samples, 229 have available progression-free survival information were included in the survival analysis. We obtained clinical trial data from a cohort of 229 patients who received chemotherapy (R-CHOP or CHOP-like chemotherapy). For this analysis, we predicted patient response to acalabrutinib combination using our pre-trained PAIRWISE model. We considered a tumor as BTKiCombo(+) or BTKiCombo(-) based on the similarity of PAIRWISE prediction probabilities by hierarchical clustering analysis **(Fig. 6a**). BTKiCombo(+) tumors demonstrated significantly longer progression-free survival (PFS) than BTKiCombo(-) tumors (62.4 versus 21.6 months, *P* = 0.004, the log-rank test, **Fig. 6b**). Previous studies demonstrated high confidence in the COO classification of DLBCL (Wright et al., 2003). Similarly, we sought to identify genes that were most discriminative in their expression between BTKiCombo(+) and BTKiCombo(-) DLBCLs and developed a linear predictor scores (LPS) algorithm for BTKiCombo-classification (**Methods, Supplementary Fig. 7-8**). The final LPS algorithm incorporated 50 genes and correctly classified 0.733 of the training set tumors into the subgroup to which they had been assigned by hierarchical clustering. The reproducibility of the LPS algorithm was demonstrated by its ability to correctly classify 0.704 (95% CI = 0.611-0.786) of the tumors in the validation set (**Fig. 7a, Supplementary Table. 5**). Our LPS approach identifies a transcriptome signature underlying drug synergy pattern defined by the PAIRWISE model.

**Fig. 6:**
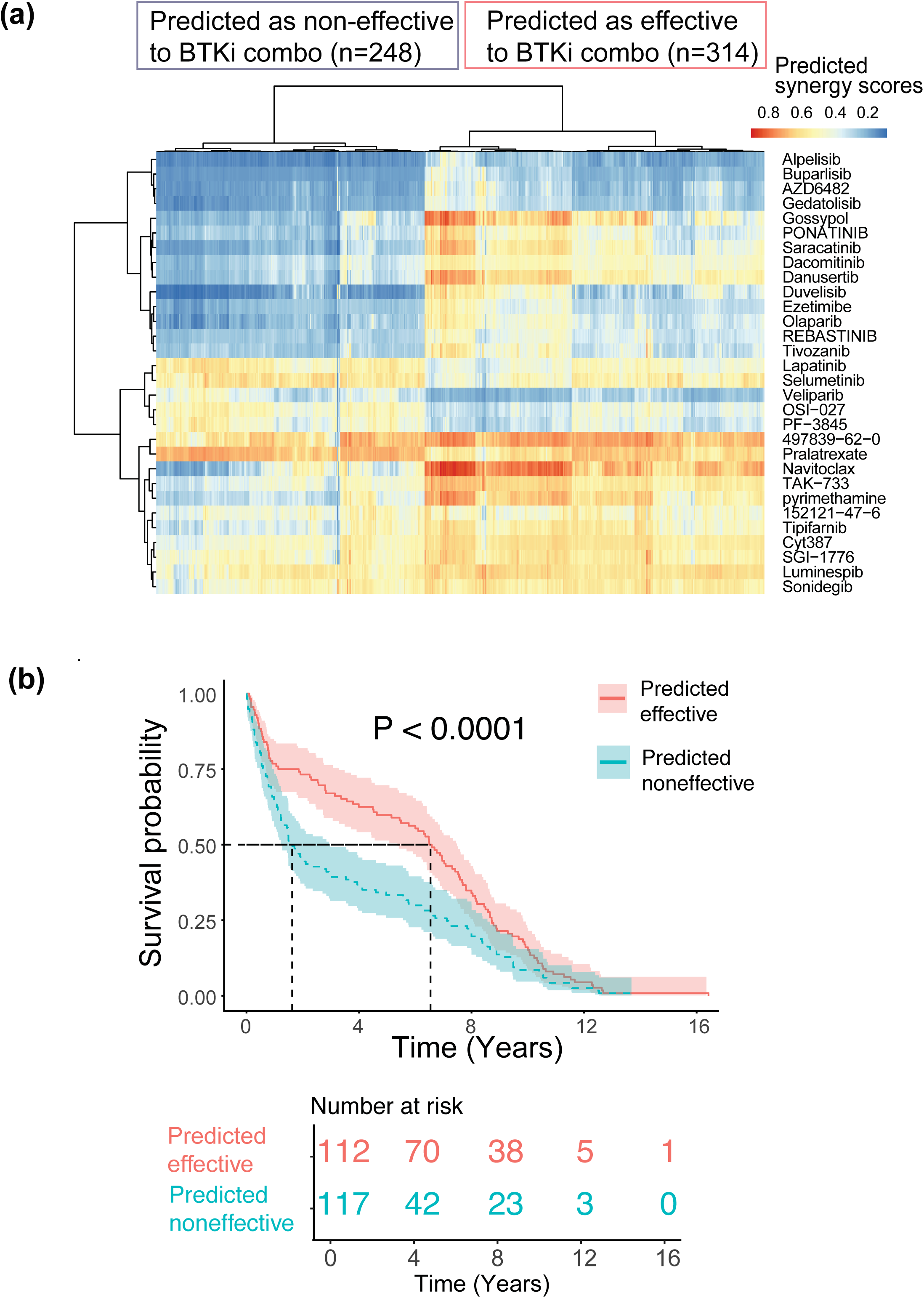
Guiding BTKi combination therapy in DLBCL patients. **(a)** The assignment of the DLBCL cases to the predicted effectiveness to BTKi combination based on the hierarchical clustering of predicted synergy scores by PAIRWISE. **(b)** Survival curves for BTKi drug combinations predicted to be effective by our model showing a significant improvement in progression-free survival. The p-value indicates significance by log-rank test.

**Fig. 7:**
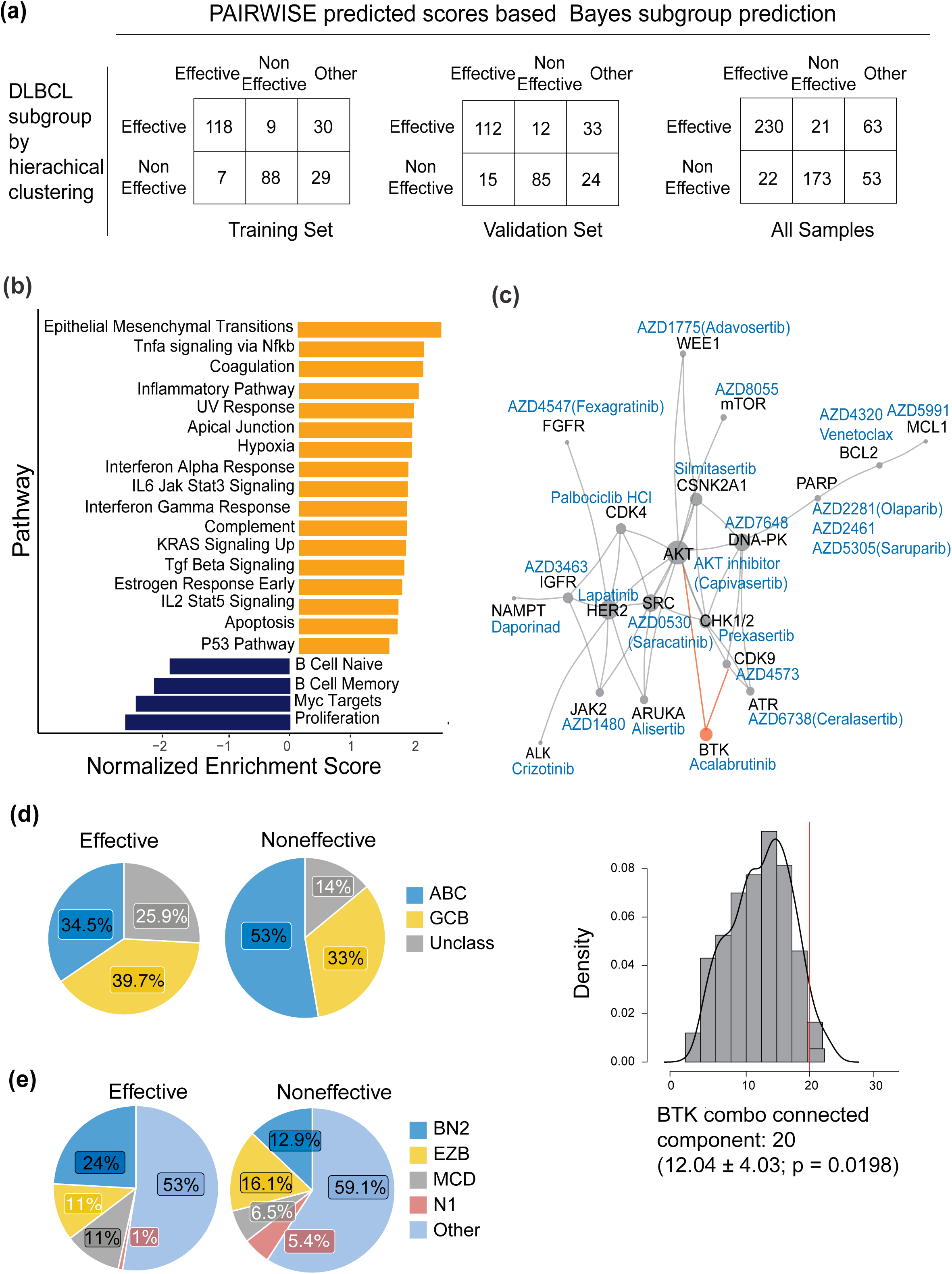
Interpretation of BTKi combination mechanisms in DLBCL patients. **(a)** The subgroup Bayes predictor are compared within the training, validation, and total set of samples. **(b)** Biological pathways most highly differentiated between DLBCL subgroups which are predicted to be effective or non-effective to BTKi combination. Pathways are sorted by their average ranking using GSEA analysis. **(c)** Network of proteins targeted by our screening BTK combo drugs (highlighted by PAIRWISE). These proteins are not distributed randomly in the human interactome, but form a connected component and multiple subgraphs shown in the graph. The bar plot shows the distribution of random expectations of the connected component size. The observed BTK combo component, whose size is indicated by the red arrow, is larger than expected by chance. **(d-e)** Distribution of predicted subgroups overlaps within gene-expression subgroups (ABC, GCB or other), or genetic subtypes (BH2, EZB, MCD, N1 or other). > 68% of younger MCD patients are predicted to be effective to BTKi combination.

Although attributions and trends for individual genes are informative, to gain system-level insights into regulatory pathways that are important to BTKi combination prediction, we applied pathway analysis (**Methods**) to determine several pathways that are differentially represented between BTKiCombo(+) and BTKiCombo(-) DLBCL. When we perform Gene Set Enrichment Analysis (GSEA) on the entire ranking of genes, we find pathways covering a broad spectrum of cellular processes and functions (**Fig. 7b**). Some of them are portions of well-known cancer pathways such as mTOR signaling, BCL2-regulated apoptosis, and epithelial-methylation transition (EMT). EMT has previously been linked to resistance to chemotherapy. BTKiCombo (+) patients would benefit from BTKi-containing combination therapies over chemotherapy.

To search for molecular mechanisms of the top drug combination candidates that are validated experimentally, we mapped the 29 candidate drug protein targets of the predicted positive BTKi combination drugs to the human interactome, a collection of 330K binding interactions between 18,508 human proteins which were assembled from 21 public databases that compile experimentally derived protein-protein (PPI) data (**Fig. 7c**). We found that 20 of the 29 involved combinations targeting proteins form a large, connected component of the PPI graph, which indicates that the predicted top candidate drug targets aggregate in the same network vicinity. Of these, one combination targets synthetic lethal pairs of genes as defined by SynLethDB including CHK1 inhibitor prexasertib (**Supplementary Fig. 9**). We also assessed the distribution of predicted subgroups overlaps within gene-expression subgroups (ABC, GCB or other), or genetic subtypes (BH2, EZB, MCD, N1 or other) (**Fig. 7d-e**). Of note, > 68% of younger MCD patients are predicted to be effective to BTKi combination. Altogether, these results indicate that a genetically distinct spectrum within DLBCL cell lines predicts the synergy that can be achieved with specific therapy combinations. PAIRWISE predictions involving DLBCL subtypes signature may therefore be useful in guiding therapy choices in the clinic.

## Discussion

Machine learning has been used extensively to predict drug combinations based on the multiple omics data available for large panels of cancer cell lines, which are crucial to developing more robust preclinical biomarkers^11^. However, the translation of model predictions into clinical impact has been very limited owing to many factors including but not restricted to data-poor cancer types, lack of learning from clinical samples and imprecise predictions of drug combinations that have been encountered during model training. A wider, more integrative analysis, along with a model that is clinically predictive, is needed to understand drug combination response that delivers value to patients. In this study, we addressed this issue with a new DL model, PAIRWISE, that enables individualized predictions on single cancer samples.

PAIRWISE consists of three branches, chemical fingerprint branch as molecular graphs, drug target branch as the binary vector, and cancer sample branch as transcriptome profiles of individual cell lines and patient tumors. The key module fusion design of the model simulates drug combination inhibition on cells through an attention mechanism. This enables the model to generalize to novel data points and make accurate predictions based on chemical fingerprints and protein targets of drugs. PAIRWISE accurately captures drug combination responses *in vitro* and that translates to *in vivo* (for example, DLBCL patient tumors) and clinical settings. We evaluated PAIRWISE’s performance extensively on the gold standard combination data spanning over 200K drug combinations across 10 cancer types. PAIRWISE not only outperformed state-of-the-art models in the public domain but also has consistent performance across both data-rich and data-poor tissues, indicating that the model had learned generalizable and transferable knowledge from chemical structures, drug targets, and molecular profiles beyond the tissue of origin. One advantage of the current study is that our approach was applied to a dataset of drug-synergy measurements in a specific DLBCL cell population, indicating a precision medicine approach. Our *in vitro* experiments show that the extent of differentiation subgroups of DLBCL cells is an important factor for the prediction of the synergy that can be achieved with specific therapy combinations, which has potential clinical applications. The potential opportunities that could emerge from our approach could be exemplified through recent clinical studies highlighting the clinical translatability of PAIRWISE’s combination recommendations across hematological cancers– acalabrutinib plus AZD4573 (CDK9i) showed high response rates with a manageable safety profile in pretreated DLBCL patients in a dose escalation study^44^, acalabrutinib plus Venetoclax (BCL2i) significantly improved progression-free survival in 1^st^ line Chronic Lymphocytic Leukemia patients^45^, and acalabrutinib plus ceralasertib (ATRi) in relapsed or refractory aggressive Non-Hodgkin’s Lymphoma in the PRISM study discontinued as of 2022 but a combination that has not yet been fully explored in DLBCL^46^. Previous study ^33^ delineated distinctive DLBCL genetic subtypes with unique genotypic, epigenetic, and clinical attributes. The rise of Single-Cell RNA-Seq technology has illuminated DLBCL’s tumor heterogeneity. PAIRWISE predictions suggested for distinct subgroup DLBCLs with mutations involvement in the DNA damage response pathway, inhibitions to DDR and B Cell Receptor (BCR) pathways can be considered for potential therapeutic responses in DLBCL.

Future work could address some of the ‘out-of-scope’ aspects of our study. Integrating mutations with additional levels of molecular information such as epigenetic states, transcriptome, or microenvironmental influences could amplify the actionable combination space. This integration could be accomplished by pre-processing multiple layers of information to derive a profile of gene scores for each cell line or tumor, which would then be input to PAIRWISE. Alternatively, to understand which information plays the most important role in the PAIRWISE model, further work can incorporate SHAP (SHapley Additive exPlanations) analysis to visualize the contribution of all the features in the best-performing. Incorporating features showing the most significant SHAP importance for the drug combination predictions can be optimally combined with PAIRWISE, which might direct the choice of incorporating biological knowledge into the modeling process. As more studies are performed measuring the specific effects of anticancer drugs on the heterogeneous individual cells and subpopulations comprising DLBCL, PAIRWISE could be applied to these datasets to rapidly scale and identify novel combination and disease mechanism insights resulting in reducing the bench-to-bedside timelines for patients.

## Methods

### Curation of the gold standard drug combination training dataset and multi-modal data for feature extraction

To train the model, we harmonized data from five large drug combination screening resources: the NCI ALMANAC, ONEIL, MIT-MELANOMA, CLOUD, and the ASTRAZENECA-DREAM dataset, as well as eight relatively small drug combination screening datasets (**Supplementary Table. 1)**. To obtain a sufficiently large drug combination dataset for model training, we curated the dataset for the uniform compound and cell line name entries. Among these data, triplets that are measured in different studies were compared and duplicate records where their synergy scores were averaged in the training dataset to reduce model over-fitting. The synergy scores (Loewe, Bliss, ZIP and HSA) were preprocessed and recomputed using the R package SynergyFinder v3.0. The gold standard drug combination screening data consisted of 279,452 cell line-drug combination triplets, covering 2,614 drug and 167 cell lines, termed the p13 dataset. All major tissue types were represented, with lung lineages the most prevalent.

To standardize drug chemical structures across datasets, we queried the PubChem entry via PUG REST for each drug’ name used in the p13 dataset to obtain an isomeric SMILES notation based on the drug name or InChIKey. Drugs with no matches in the initial search were manually annotated. In order to harmonize across multi-modal features including drug targets and their chemical fingerprints, we intended to use DrugBank ID as a unique identifier. Since there is no mapping dictionary between PubChem ID and DrugBank ID, we calculated a structural similarity profile of individual drugs based on one-hot vector representation (**Supplementary Fig 1.**). The structural similarity profile contains pairwise structural similarity scores between the input drug and all the available drugs from DrugBank which serves as an anchored comparison target. The similarity profile is generated for each drug present in the p13 dataset. Structural similarity between two drugs is measured by the Tanimoto index, which is defined as the number of common shared chemical fingerprints divided by the total number of the union of chemical fingerprints of the two drugs being compared. Beyond drug, we sought to standardize cell line name and its synonyms by obtaining its unique DepMap ID. We customized Python code via Celluorous API to retrieve a Celluorous research resource identifier (RRID) foreach cell line name appearing in the p13 dataset. We transformed RRID to a static primary key assigned by DepMap to each cell line with a mapping file ‘Sample_info_v2.csv ‘, downloaded from DepMap Public 21Q3.

In a machine learning paradigm, feature selection is essential to guarantee successful model building since the goal is to learn meaningful intrinsic relationships between input features and output targets. For predicting synergy among anti-cancer drug combinations in cell lines, input consists of both drug and cell line features that are predictive of drug synergy. Therefore we designed three branches to meet the flexibility of machine learning model inputs, namely chemical fingerprint branch, drug target branch, and cancer sample branch. In the ***chemical fingerprint branch***, PAIRWISE model takes molecular graphs of drug molecules as inputs, this was done by converting the SMILE strings into molecular graphs with a Python package RDKit (version 2019). The variant model, e.g. PAIRWISE-Morgan, integrates alternative structural information to assess model prediction ability. PAIRWISE-Morgan takes the Morgan fingerprint (radius = 2) as inputs. Morgan fingerprint decomposes each chemical structure into molecular fragments by iteratively obtaining distinct paths through each atom of the molecule. These molecular fragments were hashed into a bit vector of length 2,048 to be used for model training. In the ***drug target branch***, the drug target profile was collected from Drug Target Commons (version 2.0), which is a cloud-based platform consisting of curated experimental drug-target interaction data. The input for PAIRWISE is a 354*4098 binary matrix where binary values indicate whether a drug targets a protein. Regarding the design of the variant model, PAIRWISE-DrugBank utilized drug target profile collected from Drugbank (5.1.10), resulting in a 354*2401 binary matrix. Inspired by the Transynergy model, in the variant PAIRWISE-PropagatedDrugBank model, the drug target profile of a 354*2401 binary matrix was processed with the RWR algorithm to obtain a novel network propagated drug targets profile (a 354*2401 continuous matrix). The continuous value between 0 and 1 indicates drug effects on the associated proteins that are usually in the same pathway of the drug targets. In the ***cancer sample branch,*** we obtained the pre-treatment cell line molecular signature from the Cancer Dependency (DepMap) database. The transcriptome (21Q4 Public, ‘CCLE_expression.csv’) was downloaded in the Cell Line Sample Info file (‘sample_info.csv’) from the DepMap portal. To identify a subset of potentially significant genes that contribute to drug synergy, we selected marker genes from four sources which can effectively capture the heterogeneity of different samples: a landmark gene set comprising 978 genes from the LINCS project; the top 15% most differentially expressed genes in cancer (n = 3,008) for cancer cell lines in the CCLE project; furthermore, based on PPI network contained in the STRING database, we filtered interactions with a combined score higher than 0.7 and then identified the top 1000 protein genes that exhibit the most interactions with other proteins; finally a set of drug-target genes derived from the previously described binary matrix were also included. We union the marker genes and the selection procedure yielded 4,079 genes, henceforth called ‘PAIRWISE marker genes’, and they were used in model construction.

### Neural network configuration, training and evaluation

#### PAIRWISE neural network model

We trained a multilayer neural network model to predict the synergy score of a cancer sample, e.g. a drug pair tested in a cancer cell line or a cancer patient tumor. The core of PAIRWISE modeling framework harnesses the power of the pre-training techniques to interpret and process three input features: chemical structure branch, drug target branch, and cancer sample branch. Each distinct features is initially transformed into a fixed-length vector using a modality-specific strategy, forming the initial for the feature fusion. A distinguishing strength of this framework lies in its integration of a cross-modal feature fusion module based on the transformer architecture. PAIRWISE acts to aggregate and employ a cross-attention mechanism on these vector inputs. This capability enhances the model’s robustness and predictive power, allowing it to adeptly simulate drug-cell, and drug-drug interactions, and achieve feature updating. The attention mechanism simulating the interaction of different modalities is characterized with the following equation,

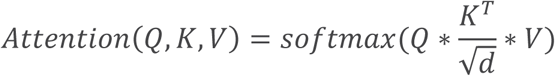

where Q(queries), K(keys), V(values) are generated matrix with the form of XW. X represents the input feature, and W is the learnable weights that linearly transform embeddings into Q,K or V. For example, to enable the module to reflect the effect of drug targets on cell PPI networks, our cross-attention module aims to find the most relevant embeddings based on attention weights between a combination of drug target embeddings, and cancer sample embedding. The updated five features (a combination of drug chemical structures, a combination of drug targets and individual cancer sample embeddings learned by PAIWISE) were flattened and concatenated to yield final embedding. The final output layer of a single neuron represents PAIRWISE predicted drug synergy probabilities, whose loss was measured against a binary synergy label. For each sample *i*, the output of the *j+1* layer 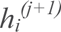 is defined as a nonlinear function of the output of the *j*th layer *h_i_*^(j)^as follows:

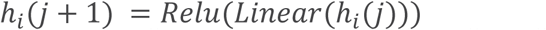

where Linear (*h_i_*^(j)^)) is linear function of *h_i_*^(j)^ defined as *W*^(j)^ *x h_i_*^(j)^ *+ b*^(j)^. *W*^(j)^ is the weight matrix and *b*^(j)^ is the bias vector. Relu is the rectified linear activation function. The first layer *h_i_*^(j)^ is the input drug chemical features, drug gene target features, and transcriptome features of sample *i* and the last layer *h_i_*^(N)^ acts as its final prediction 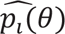, where θ is a parameter containing W(j) and b(j) from all the linear layers. In order to train PAIRWISE, we scan all the parameters and determine the architecture of the neural network by cross validation. All parameters were updated by minimizing the binary cross-entropy loss. Therefore our L term of sample set, *C*, and parameters, 0, was :

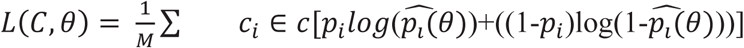

where p_i_ is the observed binary synergy label for sample *i* and *M* is the number of samples in *C.* ***Model pre-training phase for multimodal data embeddings.*** In the pre-training phase, the aim is to train individual neural network module that can quickly adapt to a new drug or a new cell line/ tumor sample with only a few additional training samples, which is often the case when a new drug pair is going to be assessed in a new sample. The rationale is to acquire prior knowledge from a set of related tasks where training samples are particularly abundant. Each modality requires a specific method of embedding. we adopted an established computational framework with “unsupervised pre-training” design of transfer learning. The transfer learning design seeks to identify universal knowledge in each modality and then to transfer this knowledge to make robust predictions in a new condition. In the chemical fingerprint branch, chemical PFM such as Grover has achieved state-of-art performance compared to other unsupervised learning techniques that were trained over millions of chemical compounds, and it is flexible such that it can be applied to drug’s 2D graph structure and output any drug’s chemical embedding. Each drug’s chemical structure was represented ‘by an average of 256 activated bits in a single vector. In the cancer sample branch, we proposed an autoencoder-based (AE-CCLE) neural network module trained over thousands of CCLE cell-line samples (n = 1,463) and TCGA tumor samples (n = 9,093) to generate biologically informative transcriptome embeddings to transfer knowledge from in vitro data into patient samples. We represented transcriptome levels by log_2_(TPM + 1), where TPM denotes the number of transcripts per million. Each cancer sample feature was represented by transcriptome data of the PAIRWISE marker gene (n = 4,079). AE-CCLE was designed with an autoencoder backbone, which was constructed to learn low-dimensional embeddings of high-dimensional transcriptomic features. We used a hyperparameter optimization method, hyperparameters, to determine the optimal number of neurons for each layer of an autoencoder. The best-performing autoencoders were then fully trained for 100 epochs. As a result, these extracted cancer sample embeddings were effectively embedded into one-dimensional vectors, each of size 256. These vectors were then combined with other modality features and directed into the prediction network of PAIRWISE for further processing. ***Model assessment and data splitting strategies.*** PAIRWISE and all comparison methods were assessed using a standard a five-fold cross-validation (CV) approach. During training, the model performance was evaluated on the validation set at the end of each epoch, and the model with the highest performance was selected. The performance of each model was measured using the area under the curve (AUROC) between actual and predicted drug synergy scores. We adopted three different methods of splitting the data for the CV folds, (1) leave-combo-out, (2) leave-drug-out, and (3) leave-sample-out. In the leave-combo-out approach, the cell line and a combination of drug triplets in the p13 dataset were divided into five folds of approximately equal size. For each fold in the cross-validation procedure, one fold of data was held out as the validation data, and the remaining four groups were pooled for training. The leave-drug-out and leave-sample-out approaches evaluate the model’s ability to predict drug combination synergy for previously unseen cell lines and compounds. This was done by splitting the drug / cell line into five clusters based on drug structure similarities / cell line transcriptome similarities. This procedure ensures that the test set did not include any drug or cell line previously present in the train set.

### Implementing benchmarked models with unified input features

Prediction performances reported by the community submitted model usually cannot be used to directly compare between models, as they typically utilized data from different sources or applied different filtering or training-validation criteria. In order to provide insightful information in terms of prediction capabilities between benchmarked models, we conducted unbiased experiments using the same p13 dataset, and the unified input feature for drug representation and/or cell line representation as previously discussed in the multi-modal dataset for feature selection. Existing studies differ mainly in terms of regarding the synergy prediction as a regression problem, by directly predicting raw synergy scores, or as a classification task, by predicting binarized labels based on pre-defined thresholds on the specific synergy scores. We selected the Loewe score (threshold for 0) as training label for comparing binary classification capabilities (synergy or non-synergy) among the benchmark models.

Previous studies showed the machine learning features that consider both drug target information and molecular profiles would improve prediction performance. We evaluated eleven representative models which utilized different feature encoders to represent different types of input features, including 5 state-of-art methods (DeepSynergy, DeepDDS, MatchMaker, MultitaskDNN, Transynergy) and 6 variant models of PAIRWISE to assess the importance of each modality. In the replacement of cancer sample branch, PAIRWISE-PathwayNN is designed to simulate the effects of drugs on cells at the pathway level to reflect the underlying signaling mechanisms. For data preparation, for each of the cancer cell lines, we computed pathway enrichment scores for 1329 canonical pathways from MSigDB. We formulated the drug combination prediction task with pathway enrichment scores, in tandem with two drugs’ descriptors as a binary classification problem. PAIRWISE-PPI model combines pharmacology networks and complex biological networks. Given a cell line graph, we used Graph Attention Network (GAN) to update gene features. Cell line graph-level representations are concatenated with drug features, and then fed to the fully connected layer to predict synergy scores.

### Independent validation of DLBCL high-throughput combinatorial screening dataset

We obtained drug screening data that identifies drugs that cooperate with ibrutinib to kill DLBCL lymphoma cells from Project Tripod from NCATS, a data-sharing platform established by the National Center for Advancing Translational Sciences. This resource contains thousands of combination pairs that can be quickly narrowed down to identify the most effective pairs for follow-up and clinical studies. We removed compounds from the dataset without known drug target or chemical structures information. Transcriptome data of TMD8 DLBCL cell line (GSE93985) was obtained from the GEO database. Such filtering left out 214 ibrutinib anchored combination assays in TMD8. We estimated the expected drug combination response using the full dose-response matrix, and the PAIRWISE classification was compared to a single synergy score.

### Translation to DLBCL patient tumors

We obtained transcriptome data and clinical information for 562 DLBCL patients. For the survival analysis, we selected all untreated patients with clinical outcome data who received immunochemotherapy (R-CHOP or CHOP-like chemotherapy; 229 samples). PAIWISE, which did not use clinical information for training, was locked down before the analysis of clinical data, allowing us to analyze the relationship between hierarchical clustered subtypes and survival in the entire cohort.

For each DLBCL patient tumor, their genomic features were used as input to our machine learning model to predict the response to BTKi that our model had not previously seen, which altogether resulted in 268 pairwise combinations with 211 drugs from diverse target classes from NCATS MIPE library and 57 additional investigational drugs. We predicted patient response to each combination using our model pre-trained on cell lines and the transcriptome profiles of each patient. We considered a combination to be “effective” if the model’s predicted drug synergy score is over 0.5. Then we performed unsupervised hierarchical clustering, which identified groups with similar characteristics. We classified a patient as BTKi Combo (+) or BTKi Combo (-) if they were in the distinct subgroups with different patterns of combination scores. We used a log-rank test (p<0.05) to determine the significance of the associated treatment outcomes (overall survival).

### Drug-specific pathway analysis

To analyze the biological processes relevant for combinations containing BTKi in the dataset, we performed differential expression analysis. Raw RNAseq count data for the BTKi Combo responsive (+) and BTKi Combo non-responsive (-) were utilized. Raw counts were transformed to log2 counts per transcripts (log-TPM) and genes with low expression levels were removed (TPM < 0.01) as previously described. Differential expression was determined by using the linear model limma. We used the set of pathways from the Molecular Signature Database (MsigDB)^47^ or the lymphoid signature database^48^for enrichment tests. We identified genes that were significantly differentially expressed when contracting the BTKi Combo (+) and BTKi Combo (-) cell lines. FDR correction was applied using the Benjamini-Hochberg procedure. Pathways enriched for these genes include mesenchymal transition and NFkB signaling, as well as pathways related with inflammatory response and KRAS signaling.

### High-throughput drug combination screening

Lymphoma cell lines including ABC-DLBCL cell lines: U2932, OCI−LY3, SUDHL−2, SUDHL−10, OCI−LY10, TMD8: GCB-DLBCL cell lines: FARAGE, SUDHL−4 were generously donated by the Melnick lab at Weill Cornell Medicine. Cell lines were cultured in basal medium [DMEM (Gibco) supplanted with 10% FBS and 1% penicillin/streptomycin (P/S)] and maintained in a 37° incubator at 5% CO2. The HTS drug-screening library composed of 30 targeted-compounds were purchased from MedchemExpress, and a subset of them were used as AstraZeneca ‘investigational drugs. Drugs were selected based on current clinical applications (FDA approved), selectivity (target of canonical signaling pathways JAK/STAT, Ras/ERK, PI3K/ATK, anti-apoptotic etc.) and redundancy (multiple drugs targeting the same pathways). Collectively a total of 21 proteins we targeted.

The high-throughput screen was conducted using a fully integrated HighRes Biosolutions automation platform, controlled by Cellario software. The system incorporated a Hamilton Microlab Star liquid handler, a Prime Automated Liquid Handler, a HighRes Biosolutions Microspin plate centrifuge, plate incubators, and a Biotek Synergy H4 plate reader.

Prior to combination screening, the single-agent activity of the drug library (MedChemExpress) was assessed to determine cell line-specific synergistic effects. Results from this single-agent screening indicated that a dose range between ∼10 nM and 10 μM would capture both active and inactive concentrations. The combination experiments utilized a 6×6 matrix block design, with customized starting concentrations and serial 1:3 dilutions for each agent, handled by the Hamilton Microlab Star.

Lymphoma cell lines were seeded in 384-well plates at a density optimized for each cell line’s proliferation, using the Prime Automated Liquid Handler. Cells were incubated for 96 hours in the presence of the drug combinations. Following incubation, the assay readout was performed using CellTiter-Glo^®^ (Promega) reagent, following the manufacturer’s protocol. Luminescence was measured using the Biotek Synergy H4 plate reader.

Quality control was ensured by monitoring the coefficient of variation (CV) of the DMSO control wells, with an acceptable CV threshold of <2%. Data analysis, including processing and visualization, was carried out using PAIRWISE Explorer.

### Key modules of benchmark platform

Since we want to evaluate which model architectures can potentially correspond well to the true biological relations between selected sets of features and ‘predictive performance’, we curated a gold-standard p13 dataset with the search space of more than 200,000 drug combinations with unified identifiers for drugs and cell lines, far beyond any individual release of dataset reported by each model in the original paper. The benchmark platform allows users to (i) load preset or customized drug combination data from a local file; (ii) specify the type of representation methods for drug and cell lines; (iii) split the dataset into training, validation and testing sets; (iv) initialize a machine learning model or load a pre-saved model with configuration files containing saved model architecture and parameters. (v) train the model and monitor the progress of training and performance metrics.

### End-to-end pipeline to compare PAIRWISE model with other models

First, we curated ground-truth datasets for both cell-line and clinically tested drug combinations. Second, we develop PAIRWISE model and re-construct other drug combination prediction models using the same drugs and transcriptome inputs. Our pipeline is capable of modeling heterogeneous data at multiple scales. Third, we benchmark PAIRWISE model with other community submitted methods related to the discovery and development of effective combination therapies. Through extensive experiments on selected datasets, we demonstrated our model has attained superior predictive performance and generalization ability to novel data points. Additionally, our pipeline offers a range of data functions including various types of molecule feature generation, strategies for systematic model evaluation, and interactive visualization for comparison and exploration, all of which are integrated and accessible through an open Python library. Last, we apply PAIRWISE model to a dataset of ex vivo anticancer drug synergy measurements for patients with DLBCL, identifying not only biological processes that are important for the determination of drug synergy but also uncovering clinical benefits on the sub-groups of patients that are useful for personalized drug combination recommendations.

We implemented an end-to-end pipeline to assist the development and evaluation of drug combination prediction models (**Supplementary Fig. 2**). Next, to benchmark PAIRWISE with state-of-the-art methods, our pipeline incorporated PAIRWISE along with 5 previously published methods predicting responses of combination treatments and 6 variant PAIRWISE models with distinct drug chemical, drug target, and cancer representations (**Supplementary Table. 2)**. All these models were developed using the same transcriptome profile or drug chemical datasets as the PAIRWISE model despite their unique feature engineering purpose. Third, our pipeline enables users for systematic evaluation and comparison among different methods using the newly curated database. The prediction results of each model can be accessed through the website (**Code availability)**. Finally, we applied PAIRWISE into real-world scenarios, with patient tumors or drug compound that were never seen before by the model.

## Code availability

PAIRWISE and the benchmarking platform can be downloaded from GitHub (https://github.com/Mew233/pairwise). Prediction results of PAIRWISE and all benchmarked models can be accessed on ShinyApp (https://mew233.shinyapps.io/synergyy_shinyr/). BTKi drug combination experimental data in DLBCL can be accessed through PAIWISE explorer (https://mew233.shinyapps.io/PAIRWISE_Explorer/).

## Acknowledgement

The project was funded by AstraZeneca. O.E. is supported by UL1TR002384, UG3CA244697, R01CA194547, P01CA214274 grants from the National Institutes of Health and LLS SCOR grants 180078-02, 7021-20, 180078-01. H.P. was sponsored by Beijing Nova Program (20220484073). The authors would like to thank David Andrews, Katy Montague, Ezo Gruyters and the AstraZeneca External R&D Academic Alliances team for their help in setting up this collaboration. The authors would also like to thank the wider collaborative teams from Cornell Medical and AstraZeneca for their support and valuable feedback right through the project.

## Supplementary figures

**Supplementary Fig. 1.**
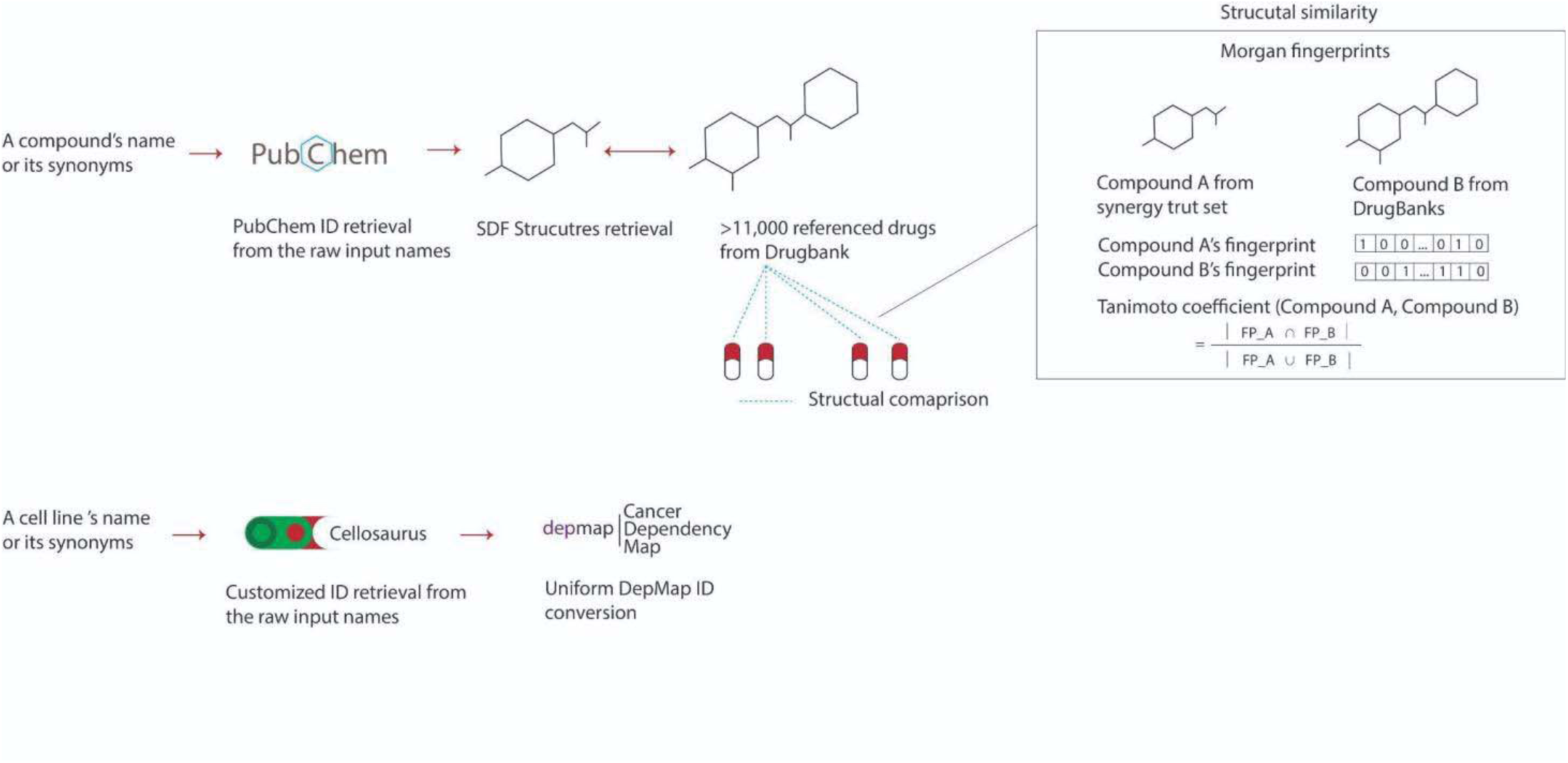
**Retrieve, harmonize, and curate drug combination datasets (pl3).**

**Supplementary Fig. 2.**
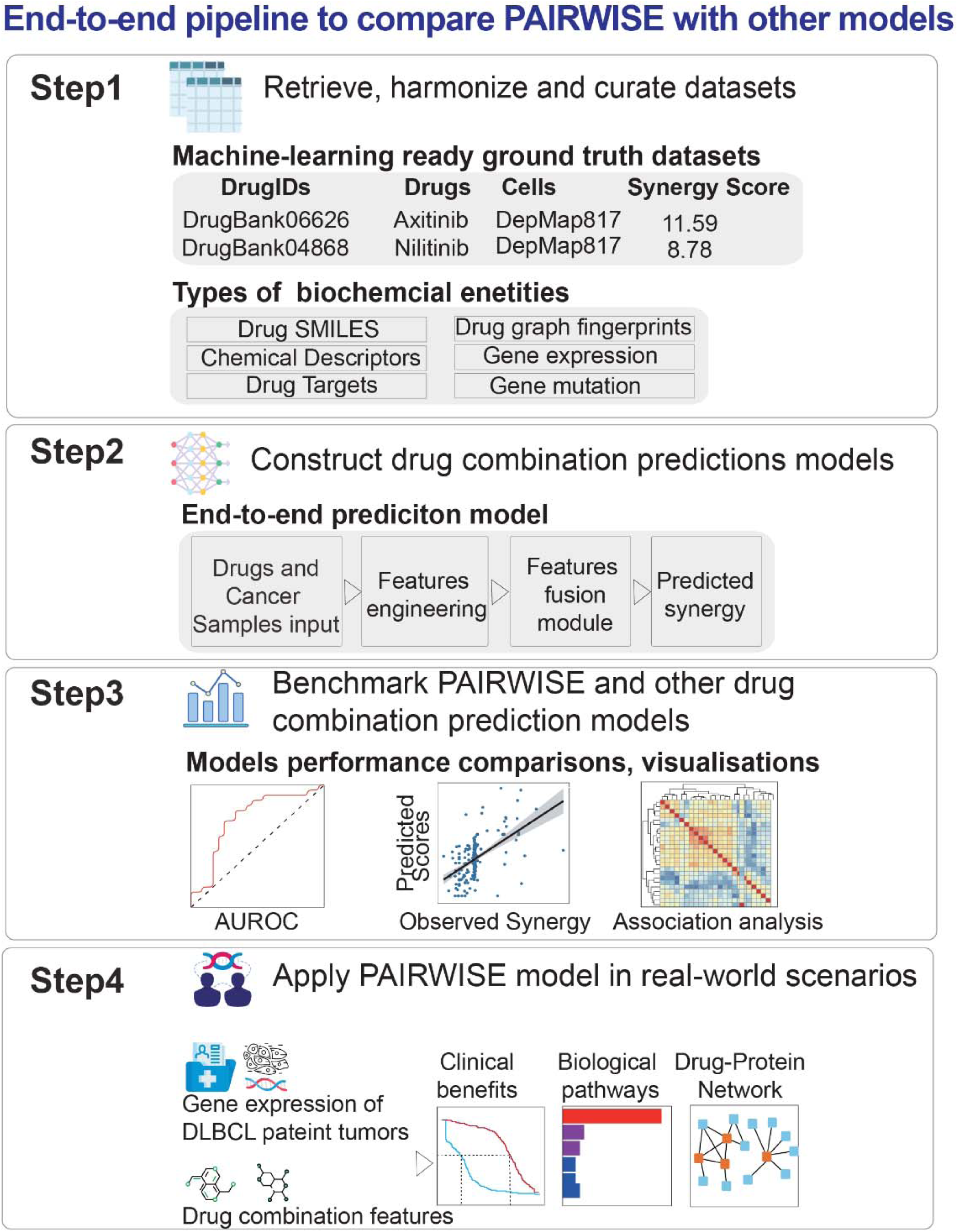
End**-to-end pipeline to compare PAIRWISE model with other models**

**Supplementary Fig. 3.**
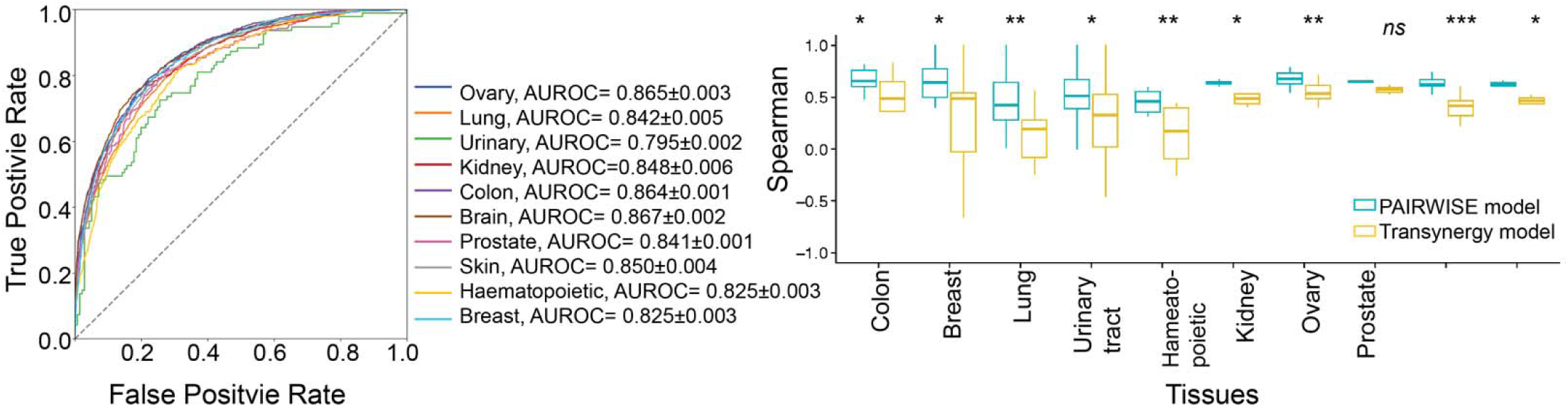
Left: PAIRWISE exhibited consistent AUROC performance across tissues. **Right:** Spearman correlation coefficients of the PAIRWISE predictions and Transynergy predictions across cell lines from different tissues of origins

**Supplementary Fig. 4.**
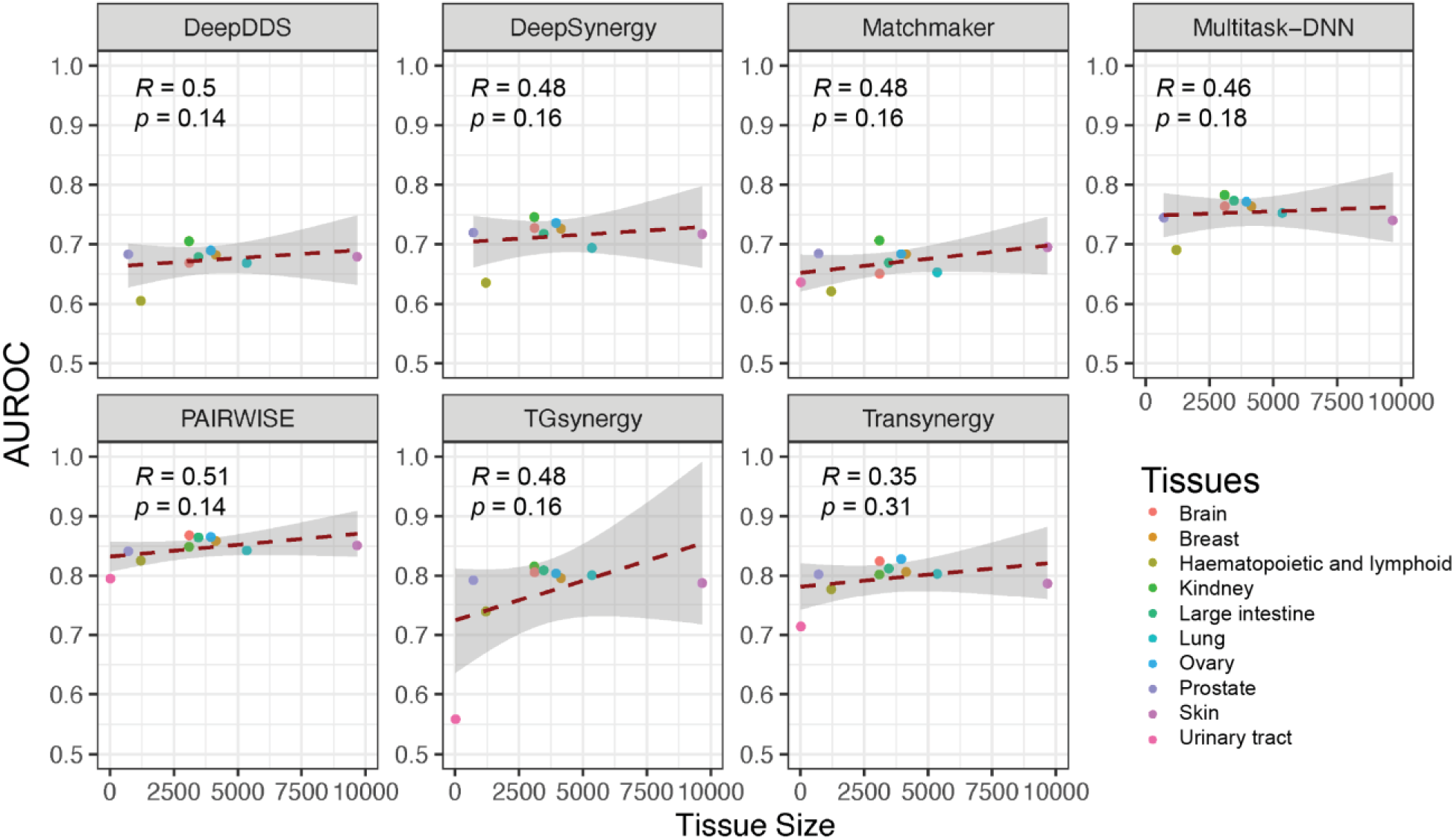
The correlation between model performance with AUROC as metrics and tissue sizes for all benchmark models.

**Supplementary Fig. 5.**
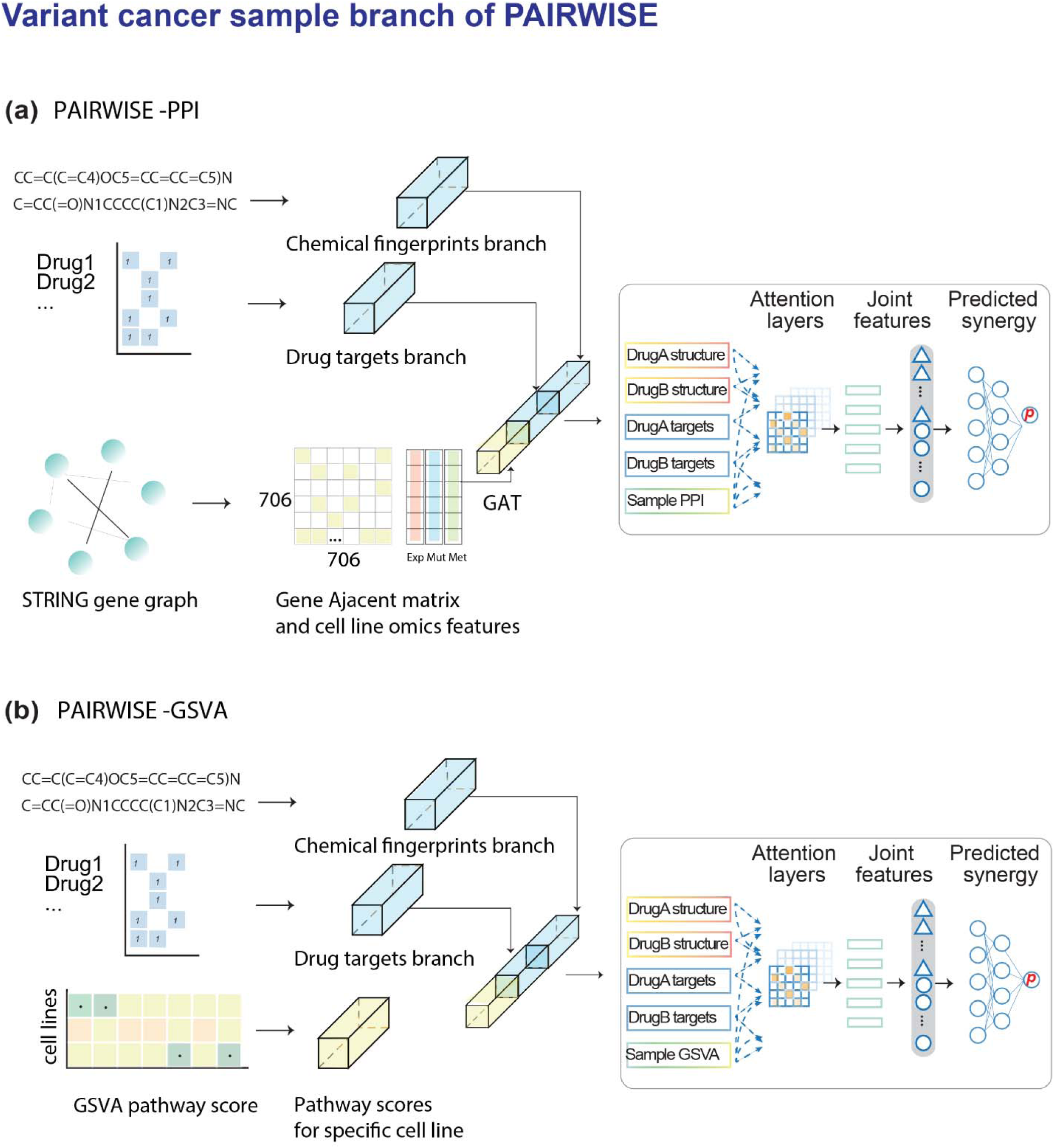
Variant models of PAIRWISE. (**a**) The cell line representation branch was derived from PPI network using a GAT structure. **(b)**The cell line representation branch was derived with GSVA pathway scores per cell line.

**Supplementary Fig. 6.**
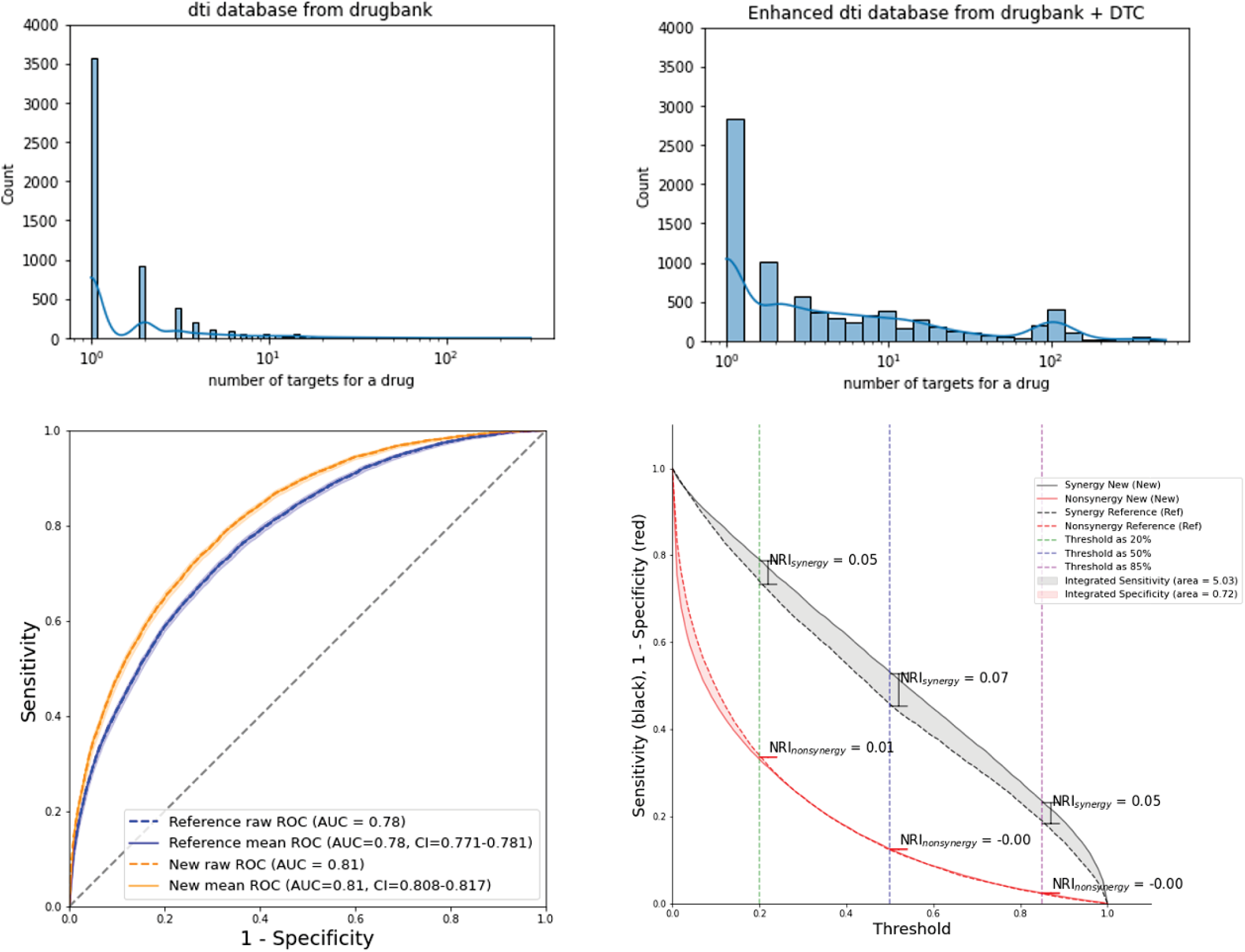
Enlarged drug-target interaction dataset enhances the predicted performance of the model.

**Supplementary Fig. 7.**
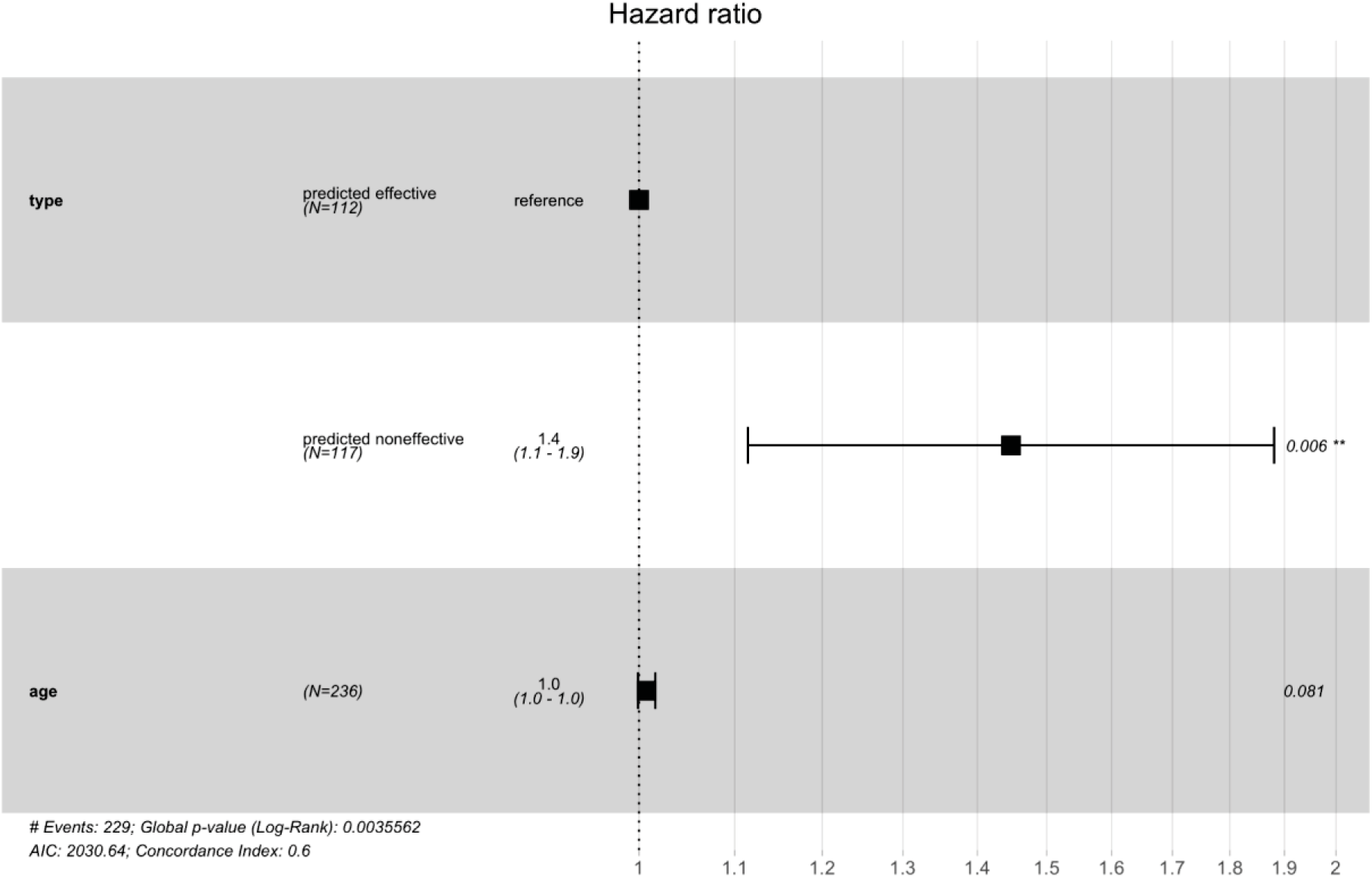
Forest plot for the BTKi-Combo (+) vs. BTKi-Combo (-) on PFS among DLBCL patients. Comparisons of the survival curves in Fig 5b were performed with a two-sided log-rank test. P values reported in this plot are two-tailed from Cox proportional hazard regression analyses. Black square represents the HR value. Error bars represent the 95% confidence intervals. PAIRWISE predicted score was associated with worse PFS in the predicted DLBCL BTKi-Combo (-) group (HR = 1.4 (1.1-1.9), P < 0.006). By contrast, age did not appear to have any effect on PFS (HR = 1 (1.0=1.0), P = 0.081).

**Supplementary Fig. 8.**
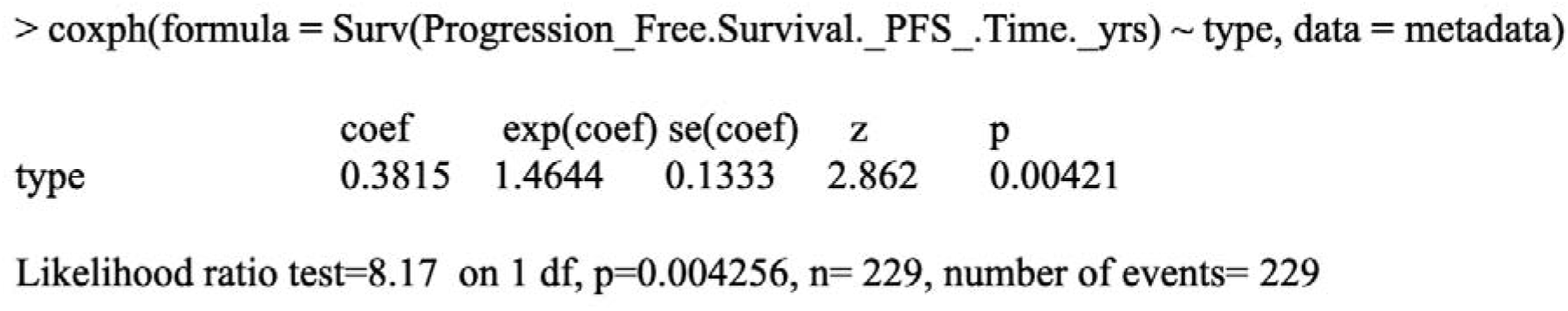
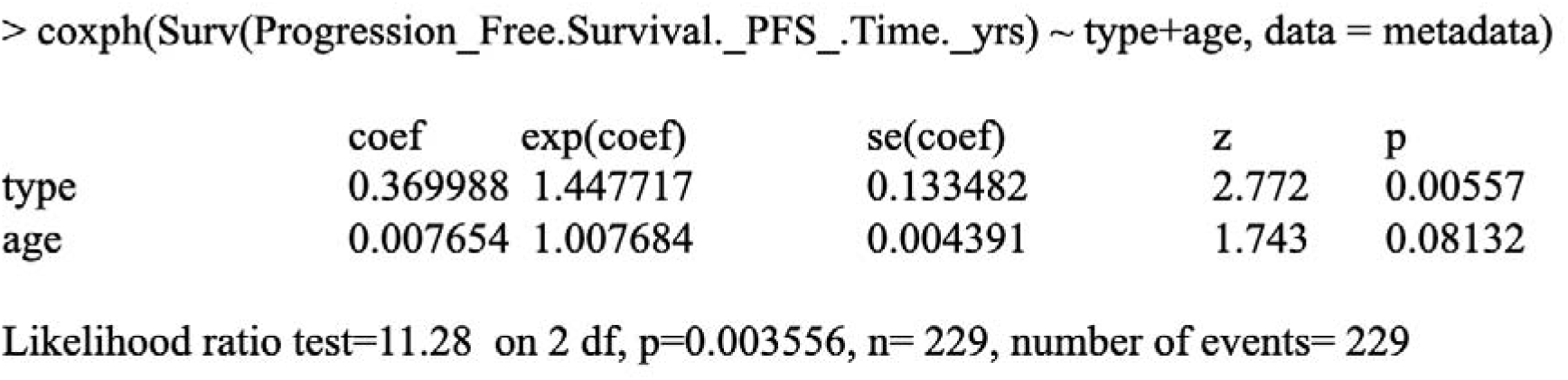
Multivariate Cox analysis for the BTKi-Combo (+) vs. BTKi-Combo (-) patients, and with consideration of covariates age. The covariate of PAIRWISE identified type remains significant (p < 0.05). However, the covariate age is not a significant contribution (coef =0.007, p = 0.08).

**Supplementary Fig. 9.**
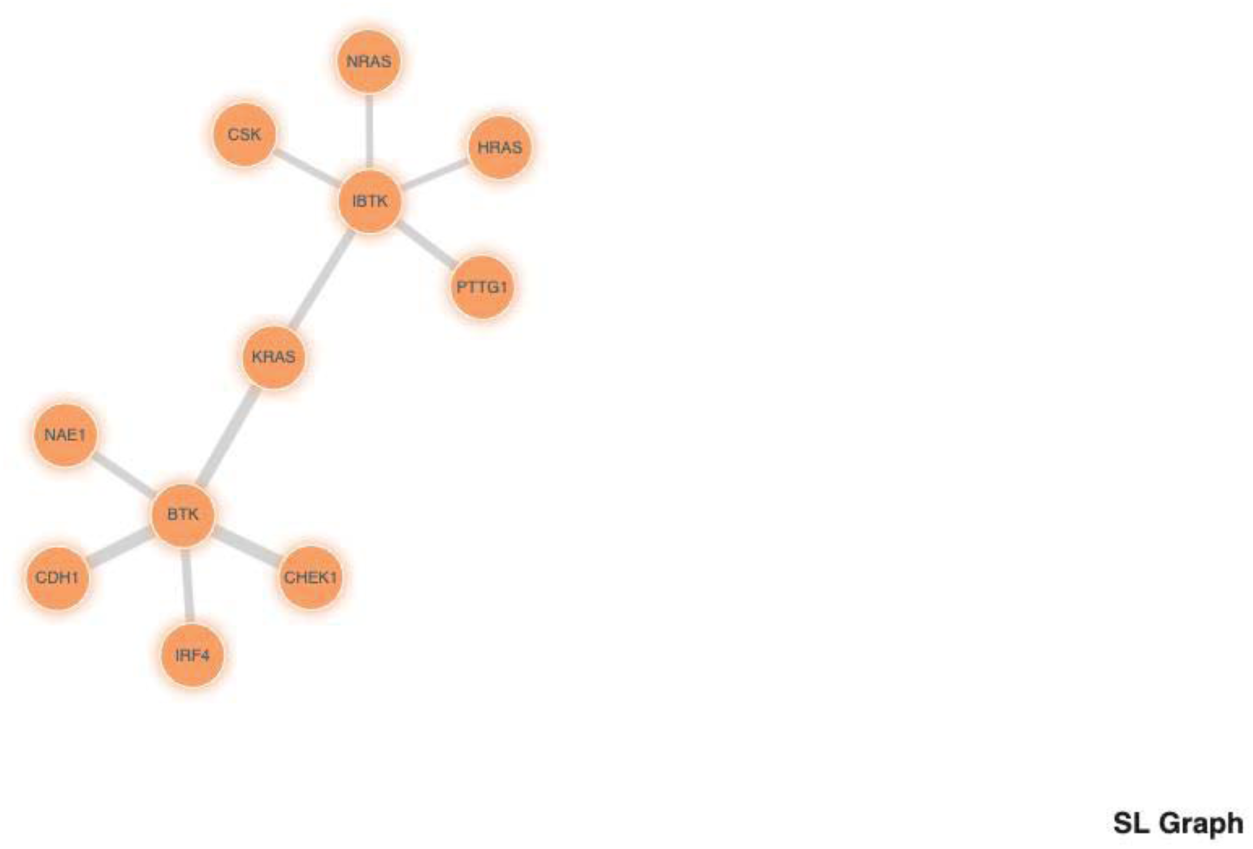
BTK combination targets synthetic lethal pairs of genes as defined by SynLethDB including CHK1, Prexasertib is such as a CHK1 inhibitor tested in our experiment.

**Supplementary Table. 1.** Characteristics of the studied datasets reported in the original paper that are used in this study. . The numbers of total experiments, screened drugs, cell lines, tissues and experimental designs (dose-response matrices) are shown below.

| Study | #experiment | #drug | #cell<br>line | #tissues | Design | Ref |
| --- | --- | --- | --- | --- | --- | --- |
| ALMANAC | 311,604 | 103 | 60 | 9 | 4-by-4 or 4-by-6 | <sup>9</sup> |
| MERCK | 92,208 | 38 | 39 | 6 | 5-by-5 | <sup>2</sup> |
| MELANOMA | 208,008 | 108 | 36 | 1 | 3-by-3 | <sup>49</sup> |
| CLOUD | 40,160 | 283 | 1 | 1 | 2-by-2 | <sup>50</sup> |
| ASTRAZENECA-DREAM | 11,576 | 118 | 85 | 10 | 6-by-6 | <sup>11</sup> |
| FLOBAK | 9,984 | 19 | 8 | 7 | 6-by-6 or 10-by-10 | <sup>6</sup> |
| YALE-TNBC | 4,576 | 130 | 6 | 1 | 1-by-5 | <sup>56</sup> |
| YALE-PDAC | 3,326 | 41 | 4 | 1 | 3-by-3 | <sup>57</sup> |
| FORCINA | 1,818 | 1,818 | 1 | 1 | 2-by-2 | <sup>51</sup> |
| GBM | 764 | 31 | 2 | 1 | 6-by-6 or 10-by-10 | <sup>53</sup> |
| YOHE | 270 | 25 | 3 | 2 | 10-by-10 | <sup>52</sup> |
| DECREASE | 210 | 33 | 13 | 1 | 8-by-8 | <sup>55</sup> |
| VISAGE | 34 | 2 | 34 | 1 | 10-by-6 | <sup>54</sup> |

**Supplementary Table. 2.**
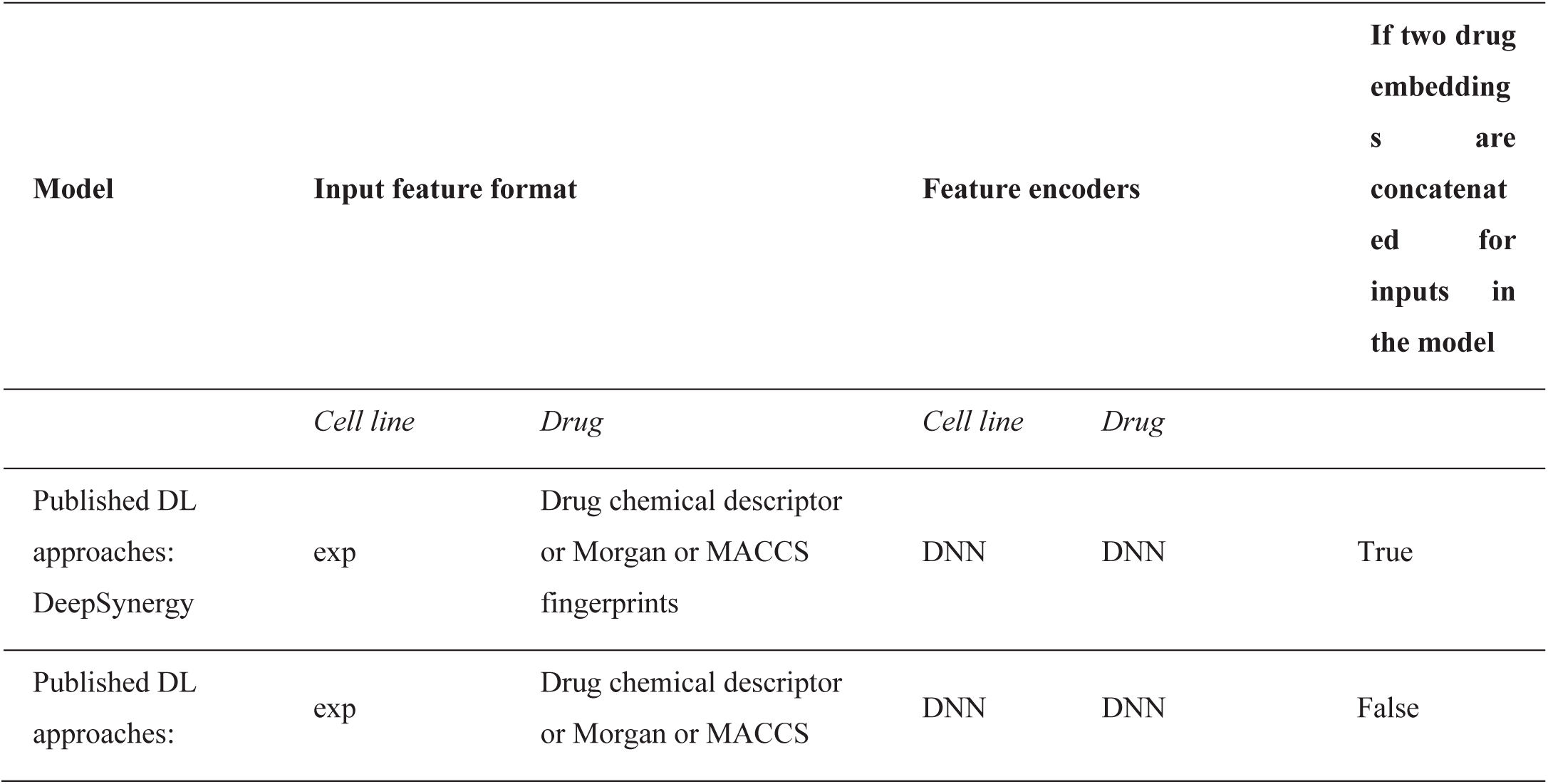
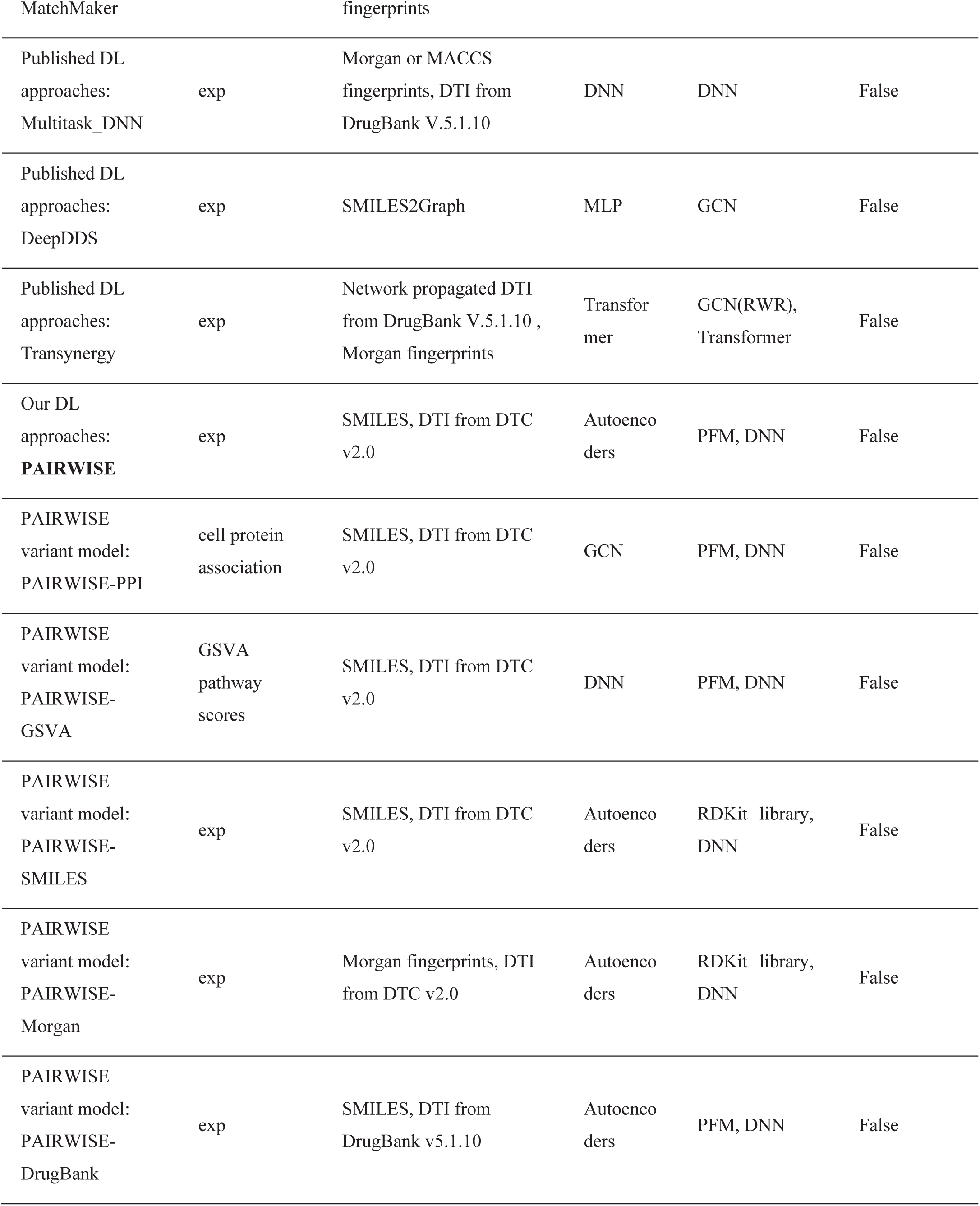
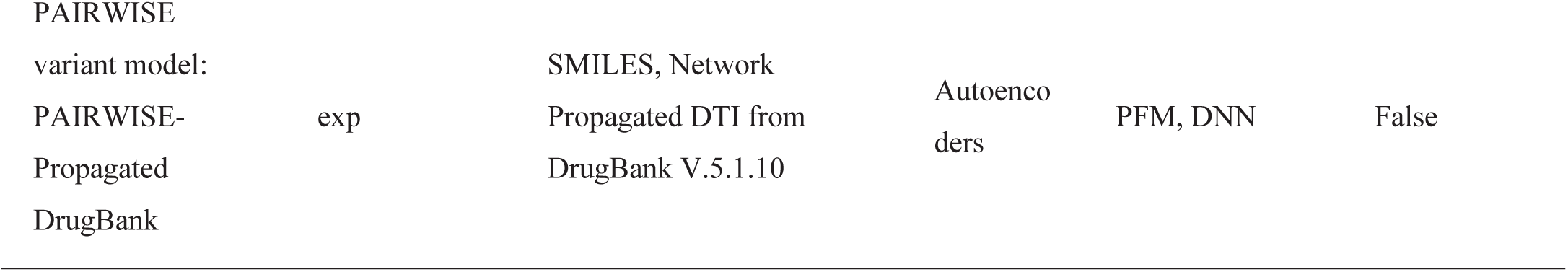
Benchmarked models for drug combination prediction in our study.

**Supplementary Table. 3.**
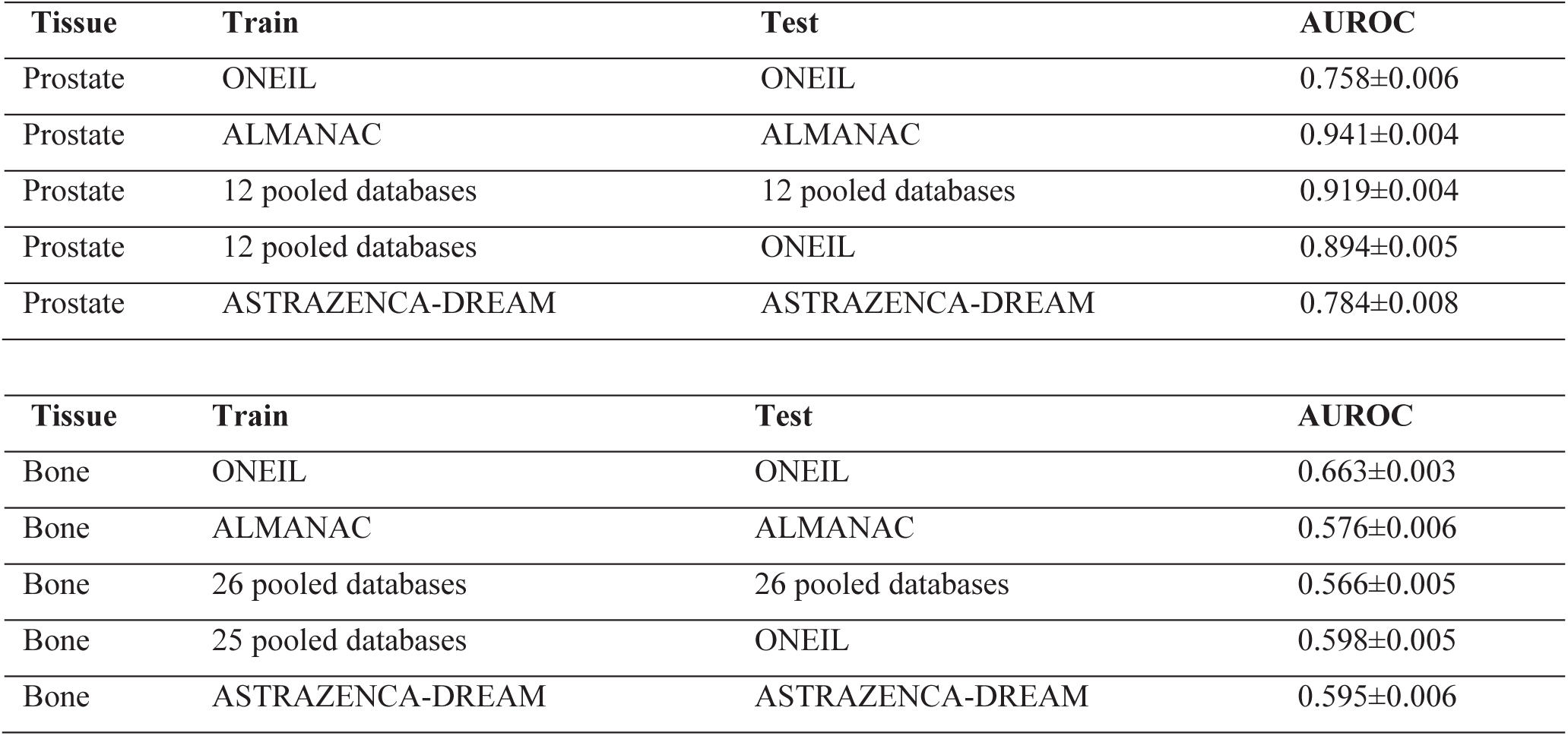
Prediction performance of specific model designed for transfer learning Multitask-DNN.

**Supplementary Table. 4 Drug screening raw data, see attachment**

**Supplementary Table. 5.** DLBCL Bayes predictor in 5-fold cross-validation.

|  | Accuracy | 95% CI |
| --- | --- | --- |
| **Fold 1** | 0.725 | (0.633, 0.805) |
| **Fold 2** | 0.651 | (0.556, 0.739) |
| **Fold 3** | 0.705 | (0.611, 0.787) |
| **Fold 4** | 0.732 | (0.640, 0.811) |
| **Fold 5** | 0.708 | (0.615, 0.789) |
| **Average** | 0.704 | (0.611,0.786) |

